# Taxonomy of *Rhizobiaceae* revisited: proposal of a new framework for genus delimitation

**DOI:** 10.1101/2021.08.02.454807

**Authors:** Nemanja Kuzmanović, Camilla Fagorzi, Alessio Mengoni, Florent Lassalle, George C diCenzo

## Abstract

The alphaproteobacterial family *Rhizobiaceae* is highly diverse, with 168 species with validly published names classified into 17 genera with validly published names. Most named genera in this family are delineated based on genomic relatedness and phylogenetic relationships, but some historically named genera show inconsistent distribution and phylogenetic breadth. Most problematic is *Rhizobium*, which is notorious for being highly paraphyletic, as most newly described species in the family being assigned to this genus without consideration for their proximity to existing genera, or the need to create novel genera. In addition, many *Rhizobiaceae* genera lack synapomorphic traits that would give them biological and ecological significance. We propose a common framework for genus delimitation within the family *Rhizobiaceae*. We propose that genera in this family should be defined as monophyletic groups in a core-genome gene phylogeny, that are separated from related species using a pairwise core-proteome average amino acid identity (cpAAI) threshold of approximately 86%. We further propose that the presence of additional genomic or phenotypic evidence can justify the division of species into separate genera even if they all share greater than 86% cpAAI. Applying this framework, we propose to reclassify *Rhizobium rhizosphaerae* and *Rhizobium oryzae* into the new genus *Xaviernesmea* gen. nov. Data is also provided to support the recently proposed genus “*Peteryoungia*”, and the reclassifications of *Rhizobium yantingense* as *Endobacterium yantingense* comb. nov., *Rhizobium petrolearium* as *Neorhizobium petrolearium* comb. nov., *Rhizobium arenae* as *Pararhizobium arenae* comb. nov., *Rhizobium tarimense* as *Pseudorhizobium tarimense* comb. nov., and *Rhizobium azooxidefex* as *Mycoplana azooxidifex* comb. nov. Lastly, we present arguments that the unification of the genera *Ensifer* and *Sinorhizobium* in Opinion 84 of the Judicial Commission is no longer justified by current genomic and phenotypic data. We thus argue that the genus *Sinorhizobium* is not illegitimate and now encompasses 17 species.

## INTRODUCTION

The family *Rhizobiaceae* of the order *Alphaproteobacteria* was proposed in 1938 and has since undergone numerous, and at times contentious, taxonomic revisions [1, 2]. Currently, this family comprises the genera *Agrobacterium, Allorhizobium, Ciceribacter, Endobacterium, Ensifer* (syn. *Sinorhizobium*), *Gellertiella, Georhizobium, Hoeflea, Lentilitoribacter, Liberibacter, Martelella, Mycoplana*, “*Neopararhizobium*”, *Neorhizobium, Pararhizobium*, “*Peteryoungia*”, *Pseudorhizobium, Rhizobium*, and *Shinella* (syn. *Crabtreella*) (https://lpsn.dsmz.de/; [3]). The *Rhizobiaceae* family contains phenotypically diverse organisms, including N2-fixing legume symbionts (known as rhizobia), plant pathogens, bacterial predators, and other soil bacteria. The agricultural and ecological significance of the family *Rhizobiaceae* has prompted the isolation and whole genome sequencing of hundreds of strains at a rate outpacing taxonomic refinement of the family. As a result, some species and genera within the family are well known to be paraphyletic [4], while others that are monophyletic likely represent multiple species/genera [5]. In addition, most currently named genera have been delineated based on genomic relatedness – as per current taxonomic guidelines [6] – but lack synapomorphic traits that would give them biological and ecological significance [7].

To aid in the taxonomic classification of this family, here we propose a general framework for defining genera in the family *Rhizobiaceae*. This framework is based on a set of baseline genomic relatedness measures meant to serve as minimal thresholds for genus demarcation, while allowing for more closely related species to be divided into separate genera when supported by supplemental genomic and/or biological data. By applying this framework, we propose the formation of a new genus – *Xaviernesmea* – on the basis of the genomic relatedness measures, and provide support for the recently proposed genus *Peteryoungia*. In addition, despite genomic relatedness values above the proposed baseline thresholds, we argue that current phylogenetic, genomic (e.g., pentanucleotide frequency), and biological (e.g., division by budding) data indicate that the genera *Ensifer* Casida *et al*. 1982 [8] and *Sinorhizobium* Chen 1988 [9] are not synonymous, meaning that the unification of the genera *Ensifer* and *Sinorhizobium* in Opinion 84 of the Judicial Commission is no longer justified.

## METHODS

### Dataset

The analysis was performed on a dataset of 94 genomes of *Rhizobiaceae* strains, among which the majority were type strains of the corresponding species (**Dataset S1**). As an outgroup, we included the genomes of three *Mesorhizobium* spp. strains, belonging to the related family *Phyllobacteriaceae*. Moreover, for calculation of some overall genome relatedness indexes to support additional taxonomic revisions, genomes of two *Pararhizobium* and two *Pseudorhizobium* strains were included (**Dataset S1**). To verify the authenticity of genomes used for taxonomic reclassifications proposed in this paper, we compared the reference 16S rRNA gene sequences (as well as housekeeping gene sequences in ambiguous cases) associated with the original species publication with the sequences retrieved from genome sequences (**Dataset S1**). Whole genome sequences (WGSs) generated to support new species description in original publications were considered as authentic (**Dataset S1**).

### Core-genome gene phylogeny

The core genome phylogeny was obtained using the GET_HOMOLOGUES software package version 10032020 [10] and the GET_PHYLOMARKERS software package Version 2.2.8_18Nov2018 [11], as described previously [12]. As a result, a set of 170 non-recombining single-copy core marker genes was selected (**Dataset S2**), and a concatenation of their codon-based alignments was used as input for IQ-Tree ModelFinder, with which a search for the best sequence evolution model was conducted. The model ‘GTR+F+ASC+R8’ was selected based on a Bayesian information criterion. The maximum likelihood (ML) core genome phylogeny was inferred under this model using IQ-TREE [13], with branch supports assessed with approximate Bayes test (-abayes) and ultrafast bootstrap with 1000 replicates (-bb 1000).

### Overall Genome Relatedness Indexes calculations

Whole-proteome amino-acid identity (wpAAI; usually simply known as AAI) was computed using the CompareM software (github.com/dparks1134/CompareM) using the aai_wf command with default parameters, i.e., ortholog identification with DIAMOND [14], e-value < 1e-3, percent identity > 30%, and alignment length > 70% the length of the protein.

Core-proteome amino-acid identity (cpAAI) was computed as the proportion of substitutions in pairwise comparisons of sequences from the 170 non-recombining, single-copy core marker genes identified using GET_PHYLOMARKERS [11], using a custom R script that notably relied on the dist.aa() function from the ‘ape’ package [15].

Percentage of conserved proteins (POCP) was determined using publicly available code (github.com/hoelzer/pocp) and the ortholog identification thresholds defined by Qin *et al*. [16], namely, e-value < 1e-5, percent identity > 40%, and alignment length > 50% the length of the protein. This pipeline involved the reannotation of genomes with Prodigal version 2.6.3 [17] and ortholog identification using the BLAST+ package, version 2.10.1 [18].

The average nucleotide identity (ANI) comparisons were conducted using PyANI version 0.2.9, with scripts employing the BLAST+ (ANIb) algorithm to align the input sequences (github.com/widdowquinn/pyani). The *in silico* DNA-DNA hybridization (isDDH) computations were performed with the Genome-to-Genome Distance Calculator (GGDC 2.1; ggdc.dsmz.de/distcalc2.php) using the recommended BLAST+ alignment and formula 2 (identities/HSP length) [19].

### 16S rRNA gene phylogeny

The RNA fasta files for the 157 *Sinorhizobium* or *Ensifer* strains analyzed in our recent study [20] were downloaded from the National Center for Biotechnology Information database, and all 16S rRNA gene sequences ≥ 1000 nt were extracted. The 16S rRNA gene sequences were aligned using MAFFT version 7.3.10 with the localpair option [21], and trimmed using trimAl version 1.4.rev22 with the automated1 option [22]. A ML phylogeny was prepared using raxmlHPC-HYBRID-AVX2 version 8.2.12 with the GTRCAT model [23]. The final phylogeny is the bootstrap best tree following 756 bootstrap replicates, as determined by the extended majority-rule consensus tree criterion.

## RESULTS AND DISCUSSION

### Overall genomic relatedness indexes measurements in the family *Rhizobiaceae*

To develop a framework for genus demarcation within the family *Rhizobiaceae*, we examined a selection of 94 genomes of *Rhizobiaceae* isolates, most of which are species type strains. *Liberibacter*, an obligate intra-cellular pathogen with a highly reduced genome, was excluded from our selection of organisms to avoid biasing the analysis by overly reducing the conserved gene set. We reasoned that good practices for genome sequence-based genus delineation should consider both phylogenetic relatedness of species based on a concatenated alignment of core-genome genes (**Figures 1, S1**), and one or more overall genomic relatedness index (OGRI) measurement. We initially considered three OGRIs, calculated as described in the Methods: (i) whole-proteome average amino acid identity (wpAAI); (ii) core-proteome average amino acid identity (cpAAI) based on the proportion of substitutions between the concatenated translated sequences of the core marker gene set used for the core-genome phylogeny; and (iii) the percentage of conserved proteins (POCP) as defined by Qin *et al*. [16]. Average nucleotide identity (ANIb; **Dataset S3**) and *in silico* DNA-DNA hybridization (isDDH; **Dataset S4**) were performed for some strains to verify they represented distinct species; however, these OGRIs were not considered when defining genera as it has been argued that these measures are more appropriate for intra-genus analyses rather than inter-genus comparisons [16].

**Figure 1.**
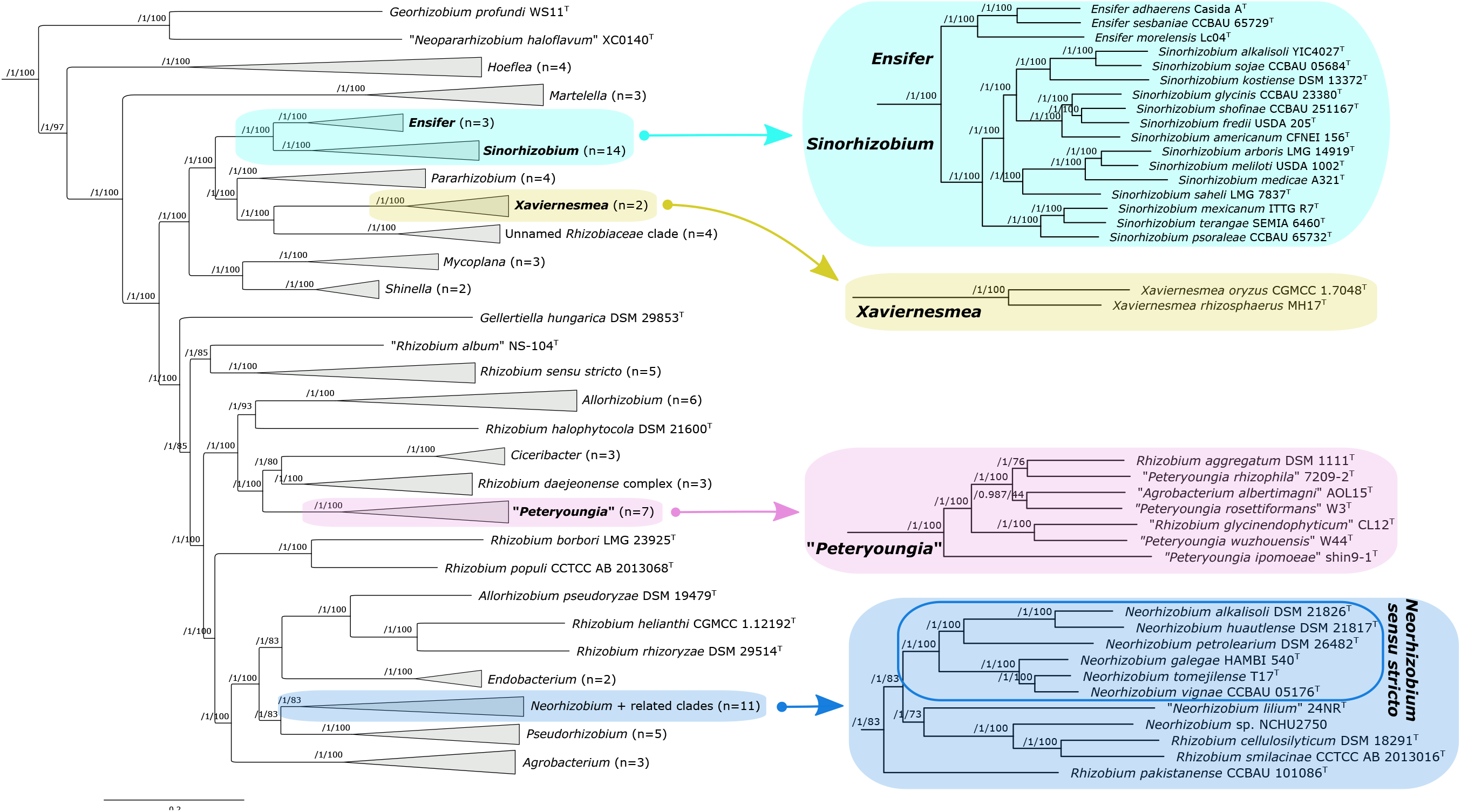
Maximum likelihood core-genome phylogeny of the family *Rhizobiaceae*. A maximum likelihood phylogeny 94 *Rhizobiaceae* strains is shown. The number of strains included in each collapsed clade is indicated. Clades of focus in the current study are expanded along the righthand side of the figure. The phylogeny is built from the concatenated alignments of 170 nonrecombinant loci using IQ-TREE [13]. The numbers on the nodes indicate the approximate Bayesian posterior probabilities support values (first value) and ultra-fast bootstrap values (second value). The tree was rooted using three *Mesorhizobium* spp. sequences as the outgroup. The scale bar represents the number of expected substitutions per site under the best-fitting GTR+F+ASC+R8 model. An expanded phylogeny is provided as Figure S1.

**Figure S1.**
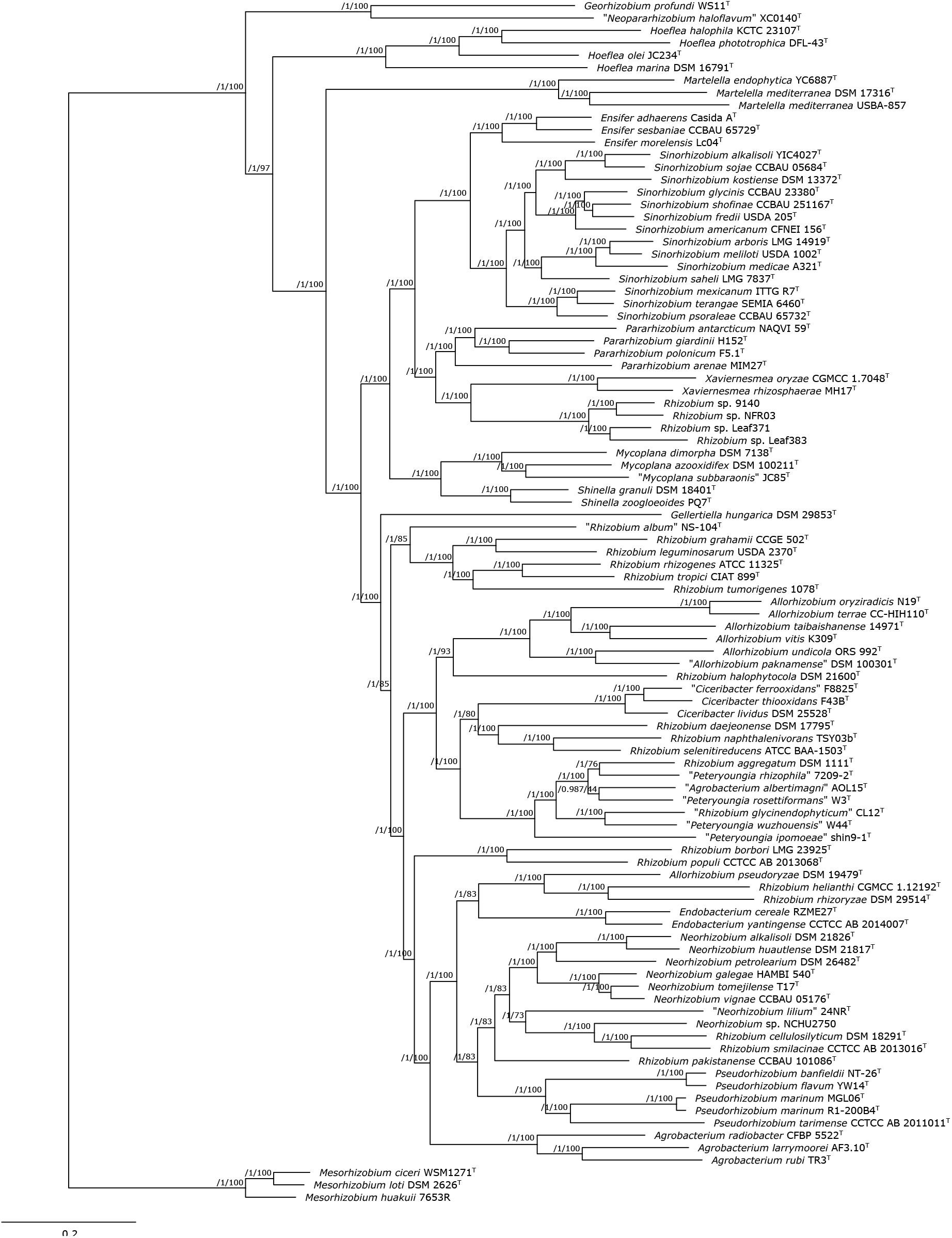
Maximum likelihood core-genome phylogeny of the family *Rhizobiaceae*. A maximum likelihood phylogeny 94 *Rhizobiaceae* strains, and three *Mesorhizobium* spp., is shown. The phylogeny is built from the concatenated alignments of 170 nonrecombinant loci using IQ-TREE. The numbers on the nodes indicate the approximate Bayesian posterior probabilities support values (first value) and ultra-fast bootstrap values (second value). The scale bar represents the number of expected substitutions per site under the best-fitting GTR+F+ASC+R8 model.

We reasoned that an OGRI threshold for delimiting genera should correspond to a drop in the OGRI frequency distribution. We therefore plotted histograms of all pairwise comparisons to identify potential genera boundaries (**Figures 2, S2, S3**). It was previously suggested that a 50% POCP threshold is a good measure of genus boundaries in other families [16]. However, we found that 3,885 out of the 4,371 (89%) pairwise comparisons in our *Rhizobiaceae* dataset gave a POCP value ≥ 50%, with no clear breaks in the frequency distribution (**Figure S2**). We therefore concluded that POCP is not a useful OGRI measurement for defining genera in the family *Rhizobiaceae*.

**Figure 2.**
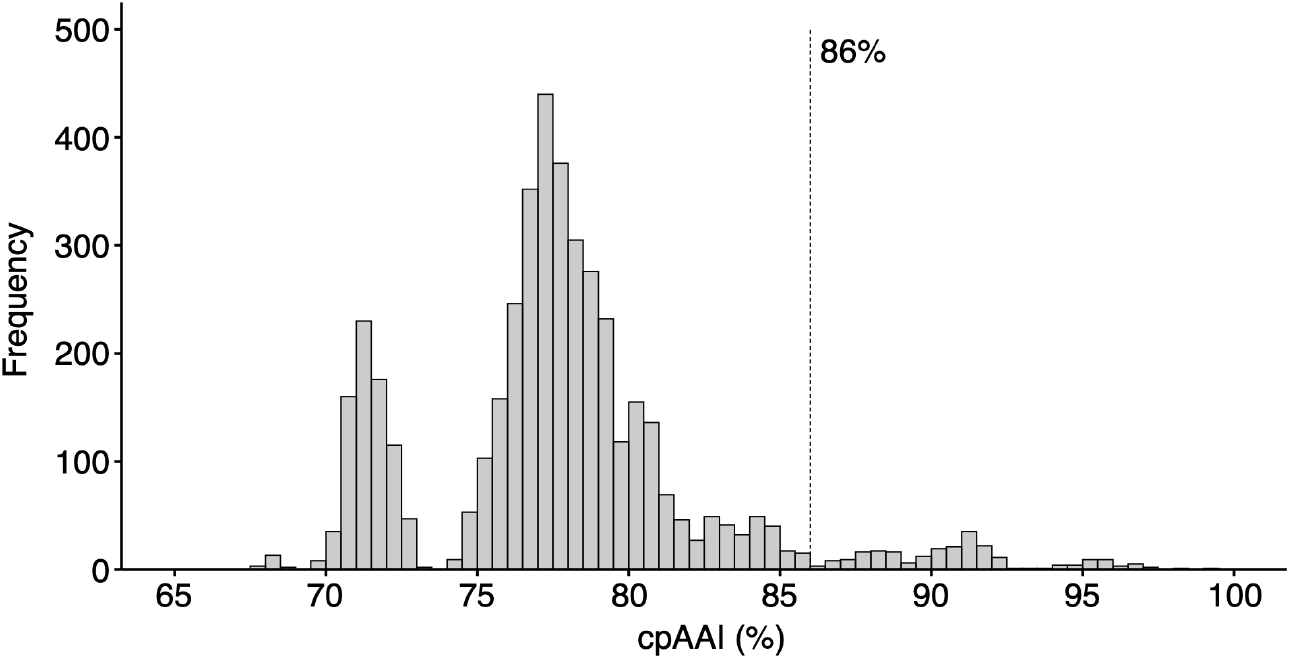
Distribution of core-proteome AAI (cpAAI) comparisons of the family *Rhizobiaceae*. Pairwise AAI values were calculated based on 170 nonrecombinant loci from the core-genome of 94 members of the family *Rhizobiaceae* Results are summarized as a histogram with a bin width of 0.5%.

**Figure S2.**
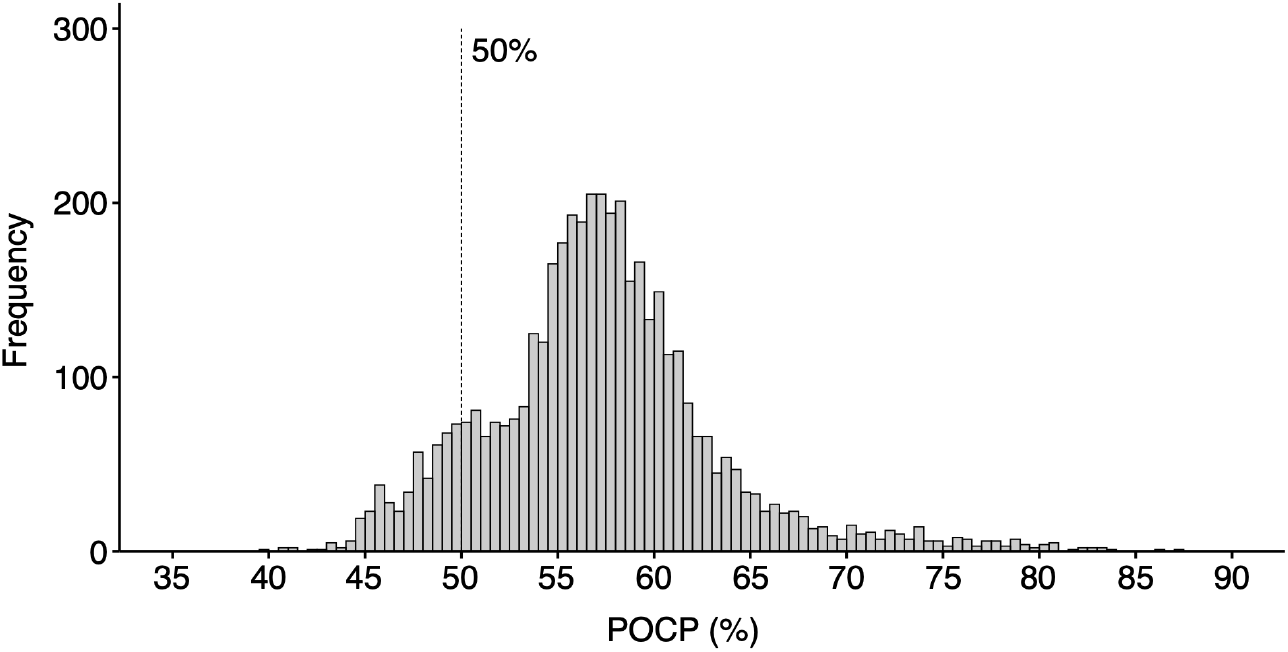
Distribution of percentage of conserved proteins (POCP) comparisons of the family *Rhizobiaceae*. Pairwise POCP values were calculated between 94 members of the family *Rhizobiaceae* as described in the Methods, and the results are summarized as a histogram with a bin width of 0.5%.

**Figure S3.**
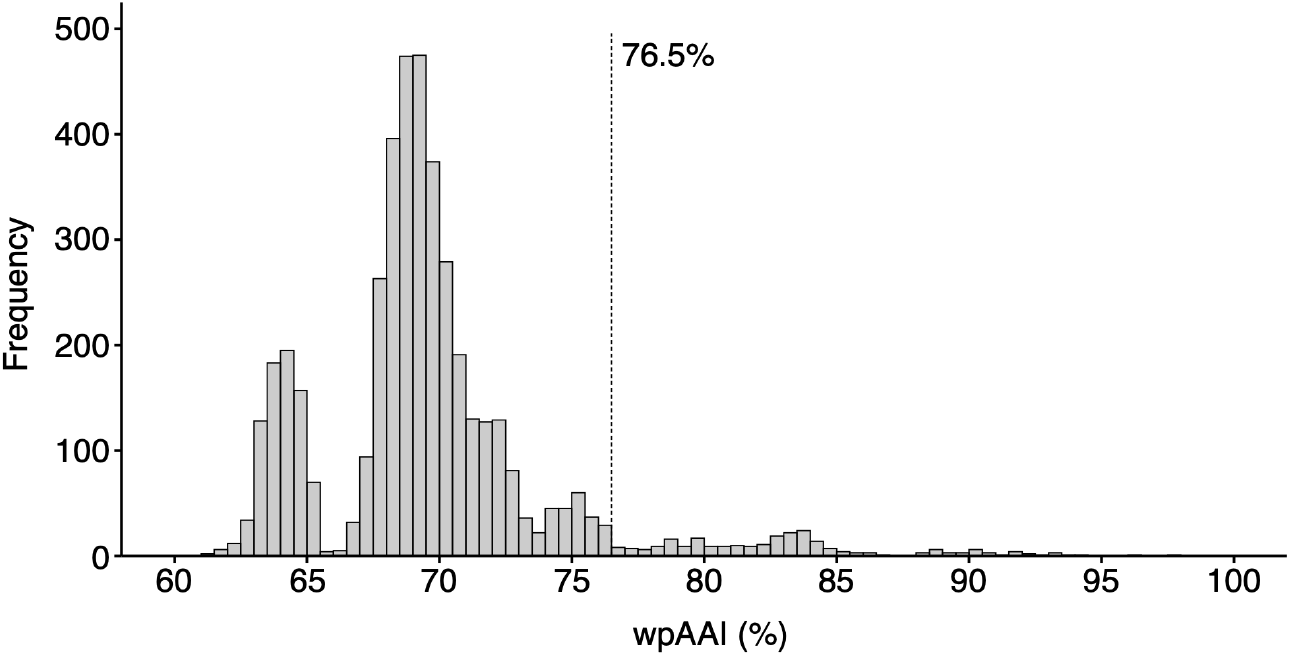
Distribution of whole-proteome AAI (wpAAI) comparisons of the family *Rhizobiaceae*. Pairwise wpAAI values were calculated between 94 members of the family *Rhizobiaceae* using the CompareM workflow, and the results are summarized as a histogram with a bin width of 0.5%

cpAAI data was recently used to delineate genera among other bacterial families [24] and stands as a promising metric for genus demarcation in the family *Rhizobiaceae* (**Figure 2**). We observed a break in the frequency distribution at ∼ 93% to ∼ 94%, but it was too stringent to use for genus demarcation as it would result in the majority of the 94 strains being classified into their own genera. Likewise, the break at ∼ 73% to ∼ 74% was too lenient for genus delimitation as all strains would be grouped as a single genus, except for those belonging to the genera *Martelella, Hoeflea*, “*Neopararhizobium*”, and *Georhizobium*. Instead, the drop in the frequency distribution at 86% to 86.7% (inclusive), within which only five of the 4,371 pairwise comparisons fell, appeared to be a reasonable threshold to aid with defining genera in the *Rhizobiaceae* family (**Figure 2**). Using a cpAAI threshold of ∼ 86%, combined with the phylogeny of **Figure 1**, we were able to recover the genera *Agrobacterium, Ciceribacter, Endobacterium, Ensifer* (as previously defined), *Gellertiella, Georhizobium, Mycoplana*, “*Neopararhizobium*”, “*Peteryoungia*”, *Pseudorhizobium, Pararhizobium*, and *Shinella* (**Figures 3, S4**). Such a threshold would, however, split the genera *Allorhizobium, Hoeflea, Martelella, Neorhizobium*, and *Rhizobium sensu stricto* into two or more genera.

**Figure 3.**
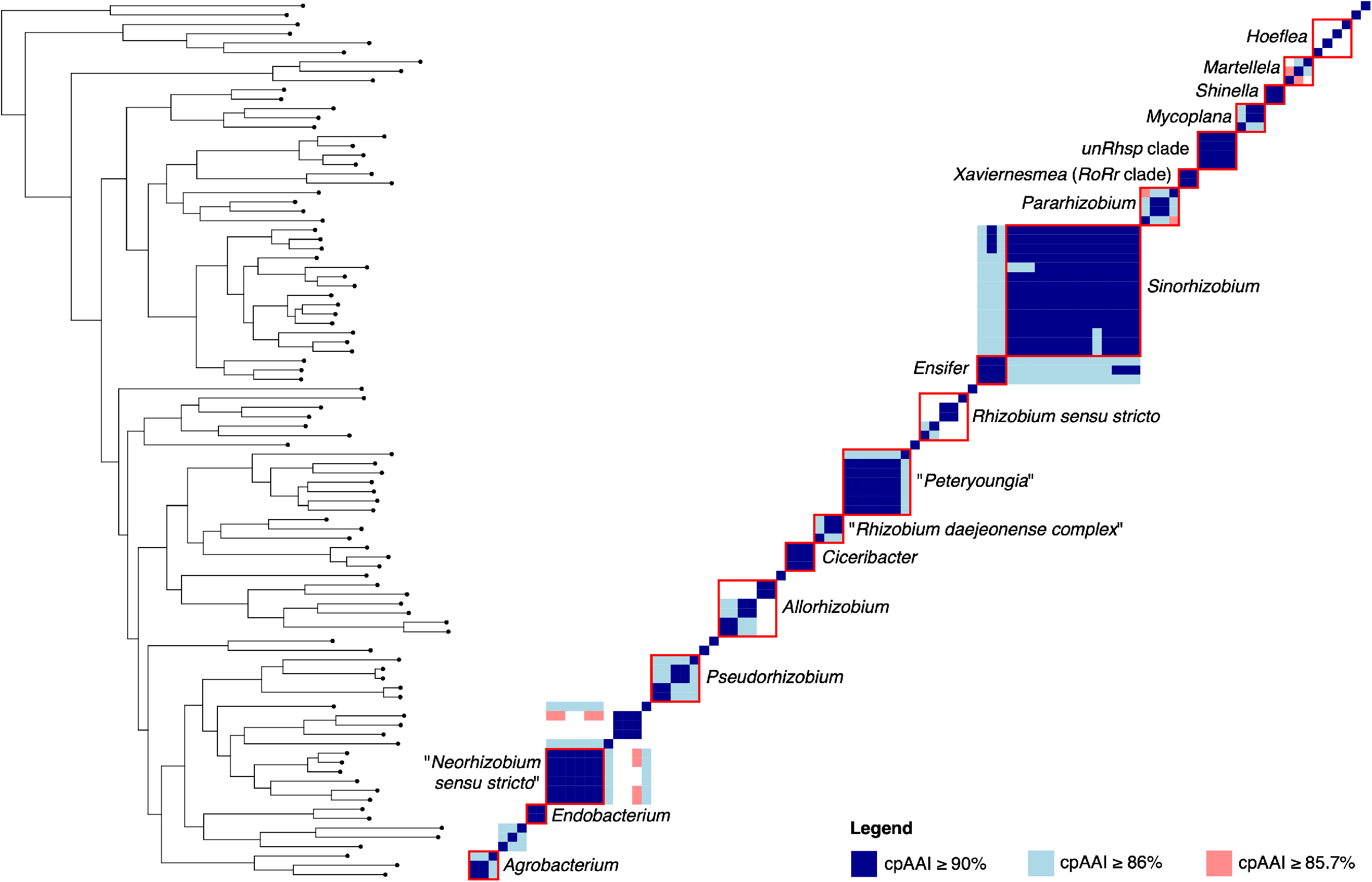
Core-proteome AAI (cpAAI) matrix of the family *Rhizobiaceae*. A matrix showing the pairwise cpAAI values for each pair of 94 members of the family *Rhizobiaceae*. Values were clustered using the core-genome gene phylogeny of Figures 1 and S1. Several named genera are indicated with red boxes, as indicated. A version of this matrix with a colour scheme representing the full range of cpAAI values is provided as Figure S4.

**Figure S4.**
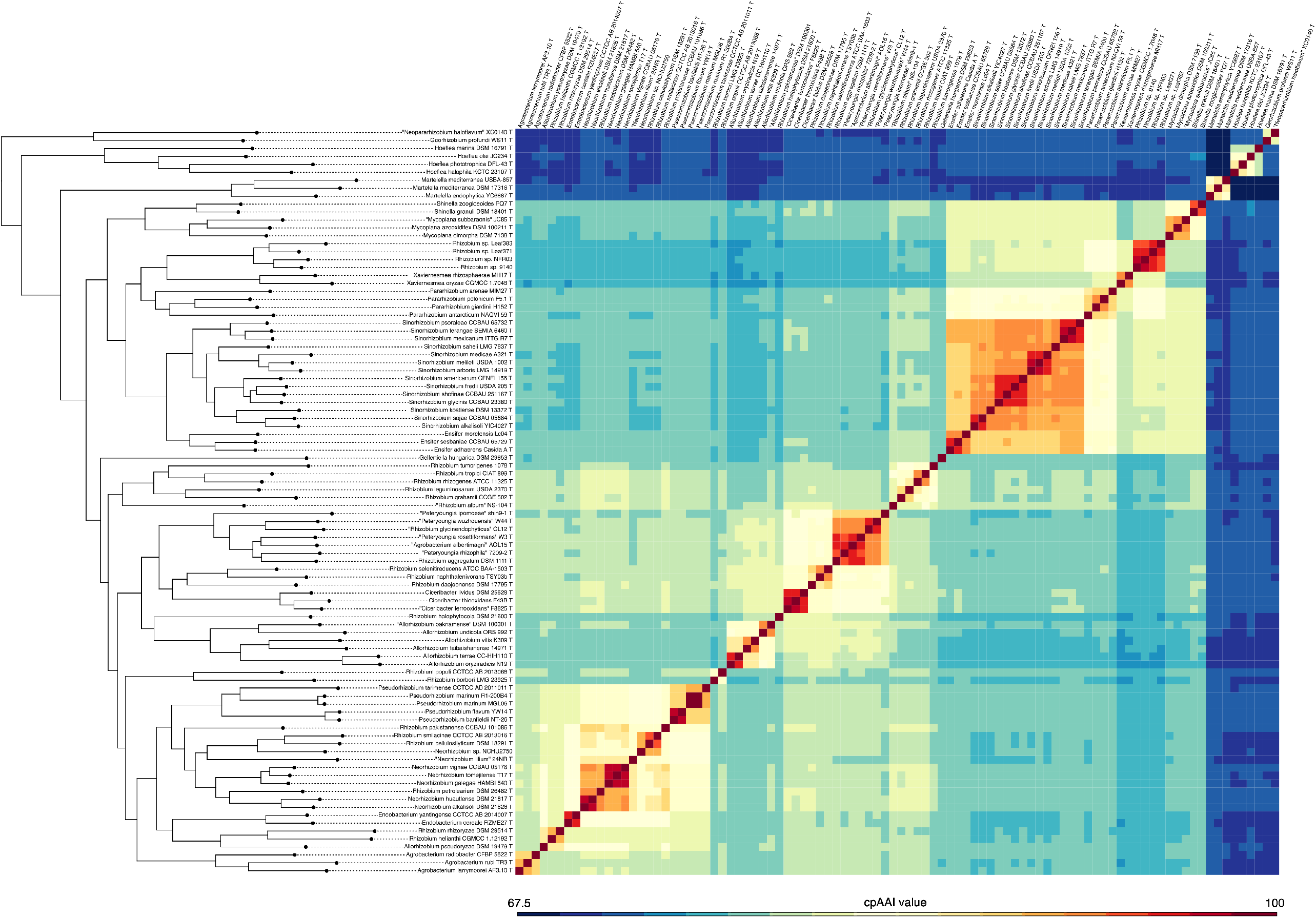
Core-proteome AAI (cpAAI) matrix of the family *Rhizobiaceae*. A matrix showing the pairwise cpAAI values for each pair of 94 members of the family *Rhizobiaceae*. Values were clustered using the core-genome gene phylogeny of Figures 1 and S1.

There was less support for the presence of a genus-level drop in the wpAAI frequency distribution (**Figure S3**). However, the wpAAI frequency distribution density increased sharply below 76.5%. Additionally, although noisier, a wpAAI threshold of 76.5% returned similar genus demarcations as did a cpAAI threshold of ∼ 86%, with the exceptions that the genus *Martelella* was recovered as a single genus, *Hoeflea* was split into fewer genera, and the separation between the genus *Ciceribacter* and its sister taxon was less clear (**Figures S5, S6**).

**Figure S5.**
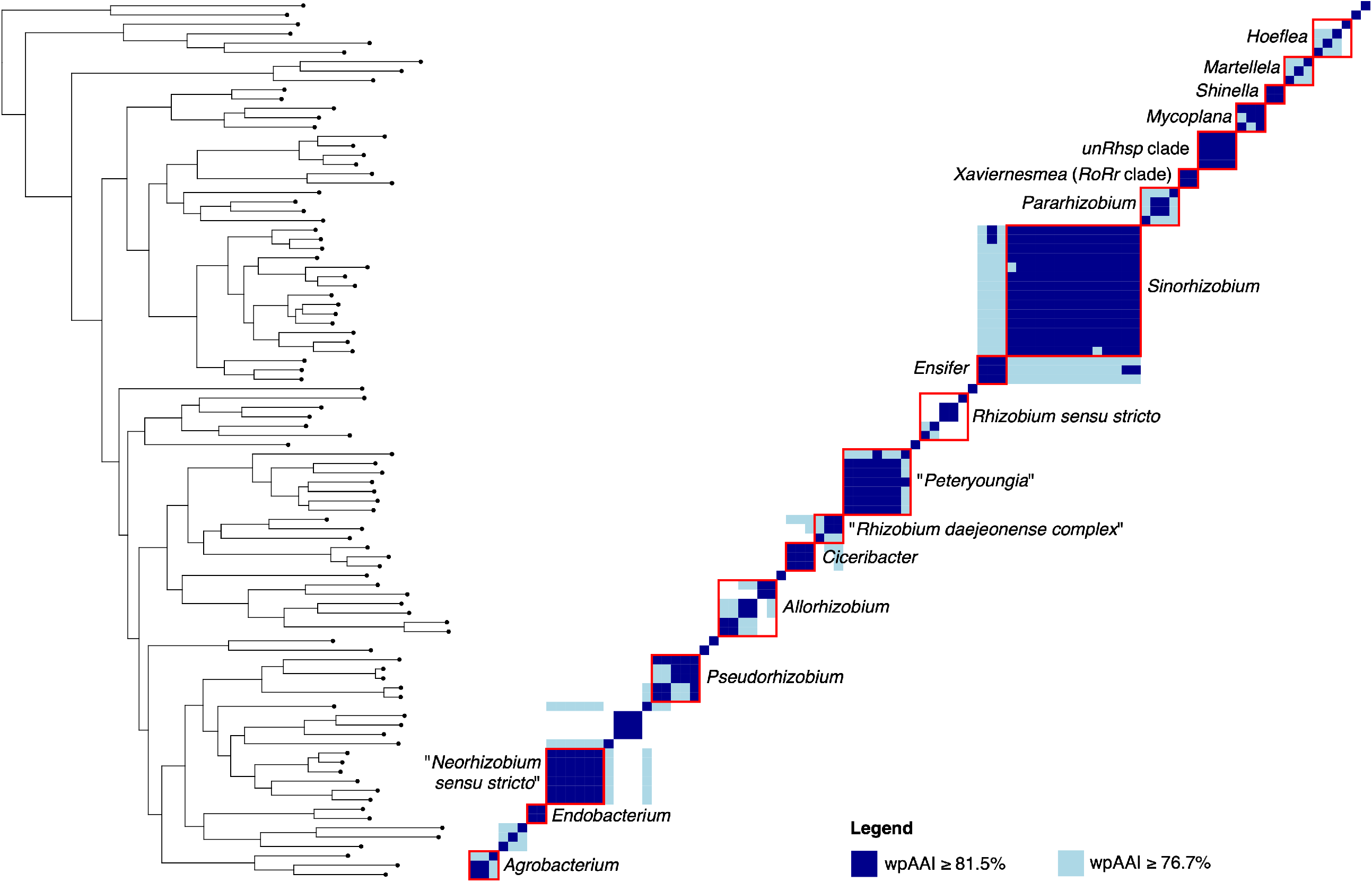
Whole-proteome AAI (wpAAI) matrix of the family *Rhizobiaceae*. A matrix showing the pairwise wpAAI values for each pair of 94 members of the family *Rhizobiaceae*. Values were clustered using the core-genome gene phylogeny of Figures 1 and S1. Several named genera are indicated with red boxes, as indicated. A version of this matrix with a colour scheme representing the full range of cpAAI values is provided as Figure S6.

**Figure S6.**
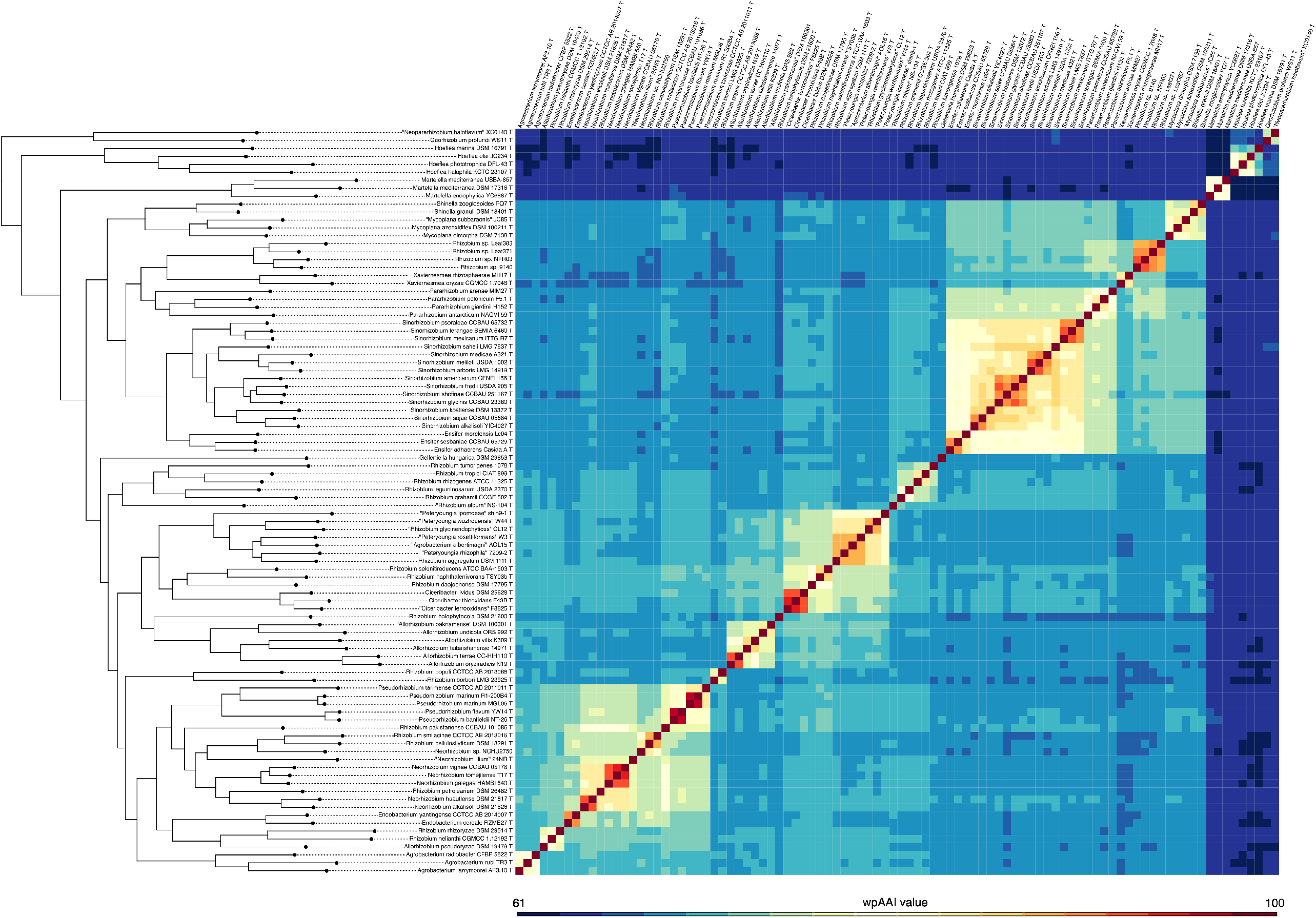
Whole-proteome AAI (wpAAI) matrix of the family *Rhizobiaceae*. A matrix showing the pairwise wpAAI values for each pair of 94 members of the family *Rhizobiaceae*. Values were clustered using the core-genome gene phylogeny of Figures 1 and S1.

### Proposal for a framework for genus delineation in the family *Rhizobiaceae*

Based on the results summarized above, we propose that genera within the family *Rhizobiaceae* be defined as monophyletic groups (as determined by a phylogenetic reconstruction using a core-genome analysis approach; **Figure 1**) separated from related species using a pairwise cpAAI threshold of approximately 86% calculated as described in the methods. We strongly recommend the use of cpAAI over wpAAI due to 1) its natural agreement – by construction – with the core-genome gene phylogeny, 2) clearer gaps in its distribution of values among *Rhizobiaceae*, and 3) the fact that it would not be sensitive to the wide genome size variation within the *Rhizobiaceae*, notably due to the variation in presence of large mobile genetic elements, including symbiotic megaplasmids. We do not, however, propose that cpAAI serve as the sole information source for genus demarcation as nearly all biological rules have exceptions. We therefore propose that genus demarcation using a cpAAI threshold higher than 86% can be justified by the presence of alternate genomic or phenotypic evidence (as proposed below for splitting of the genus *Ensifer*), while a lower cpAAI threshold may be appropriate when considering historical classifications of genera within the family.

### Taxonomic implications of the proposed framework

Following the criteria for genus demarcation outline above would notably lead to the formation of several new genera for species currently assigned to the genus *Rhizobium*, which is notoriously paraphyletic. They also imply that a few genera (*Neorhizobium, Allorhizobium, Martelella, Hoeflea*, and *Rhizobium sensu stricto*) may be candidates for division. We also note that there is a clear break in the distribution of cpAAI values at ∼ 73% to ∼ 74% that may represent an appropriate threshold for delimiting the family *Rhizobiaceae*. If adopted, this threshold would result in the genera *Martelella* and *Hoeflea* being transferred to their own families, while the genera *Georhizobium* and “*Neopararhizobium*” would form another family. However, a proposal for family-level demarcations in the order *Rhizobiales* is outside the scope of this work.

### Proposal of a new genus encompassing the species *R. oryzae* and “*R*. *rhizosphaerae*”

In a recent study presenting a phylogeny of 571 *Rhizobiaceae* and *Aurantimonadaceae* strains (ML tree based on 155 concatenated core proteins) [25], the type strains of the species *R. oryzae* (*Allorhizobium oryzae*) [26] and “*R. rhizosphaerae*” formed a well-delineated clade (with 100% bootstrap support) that was clearly separated from the closest validly published genus type, i.e. *Parahizobium giardinii* strain H152^T^. This pattern was also evident from a ML phylogeny of 797 *Rhizobiaceae* produced in another study based on the concatenation of 120 near-universal bacterial core genes [5]. The analyses presented in the current study further support the separation of the *R. oryzae*/”*R. rhizosphaerae*” clade (*RoRr* clade; two species type strains) not only from the *Pararhizobium* clade (four species type strains), but also from a sister clade consisting solely of rhizobial strains from unnamed species (*unRhsp* clade; including *Rhizobium* sp. strains Leaf383, Leaf371, 9140 and NFR03) (**Figure 1**); all three clades in the phylogenetic tree are supported by 100% bootstrap value. All within-clade pairwise cpAAI values were above 85.7%, 91%, and 94% for the *Pararhizobium* clade, *RoRr* clade, and the unnamed *Rhizobium* clade, respectively (**Figure 3**). In contrast, all pairwise cpAAI values between the *RoRr* clade and the *Pararhizobium* or *unRhsp* clades were less than 81%, while pairwise cpAAI values between the *Pararhizobium* and *unRhsp* clades were below 83.5% (**Figure 3**). All three clades thus represent separate genera according to the criteria proposed above, and this remains true when the analysis is repeated with an expanded set of strains (**Figure S7**). We therefore propose to define a new genus encompassing the *RoRr* clade, for which we propose the name *Xaviernesmea* (see below for formal description). As no strains belonging to the *unRhsp* clade have been deposited in any international culture collection, we leave the task of describing new species and genera within this clade to others who have access to these strains.

**Figure S7.**
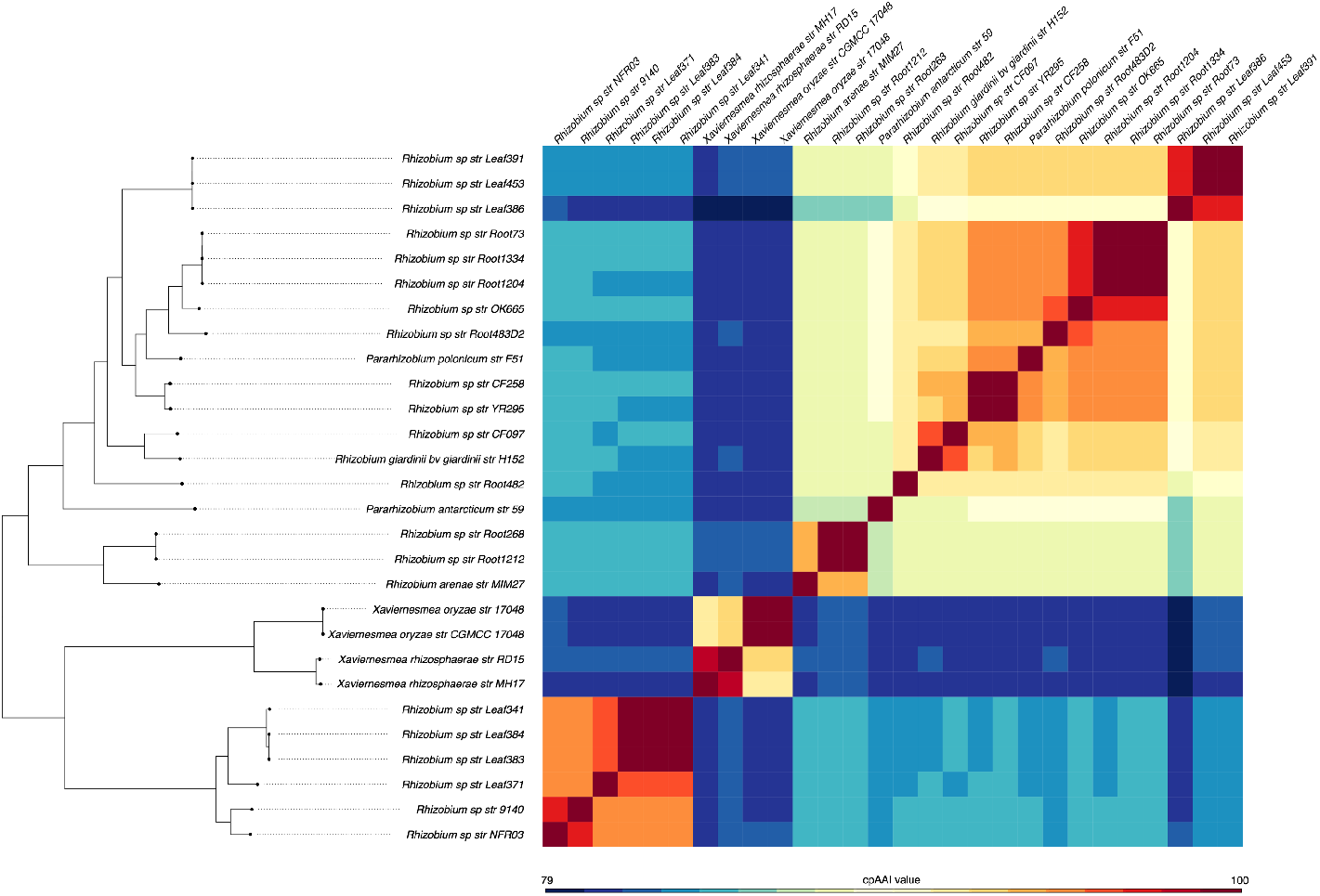
Whole-proteome AAI (wpAAI) matrix of the genus *Xaviernesmea* and related taxa. A matrix showing the pairwise wpAAI values for four members of the genus *Xaviernesmea* together with 24 related organisms.

### Taxonomy of the “*R. aggregatum* complex”

The “*Rhizobium aggregatum* complex” was initially identified as a sister taxon of the genus *Agrobacterium* [26], with subsequent work demonstrating that it is instead located on a clade neighbouring the genus *Allorhizobium* [12]. Moreover, the latter study suggested that “*R. aggregatum* complex” includes members of the genus *Ciceribacter* and that it may represent a novel genus on the basis of phylogenetic and multiple OGRI data, although the authors advised that further investigation was required [12]. It was recently suggested that the “*R. aggregatum* complex” be split into two genera [27]. It was proposed that *R. daejeonense, R. naphthalenivorans*, and *R. selenitireducens* be transferred to the genus *Ciceribacter*, while *R. ipomoeae, Rhizobium rhizophilum, R. rosettiformans, R. wuzhouense*, and “*Peteryoungia desertarenae*” be transferred to the novel genus “*Peteryoungia*” [27].

The analyses presented in the current study included 13 strains belonging to the “*R. aggregatum* complex” (**Figure 1**). The genus demarcation framework proposed here supports the previous studies indicating that the “*R. aggregatum* complex” is separate from the genus *Allorhizobium*. A group of seven species that included all *“Peteryoungia*” species present in our analysis (*Rhizobium aggregatum* DSM 1111^T^, “*Agrobacterium albertimagni*” AOL15^T^, “*Rhizobium glycinendophyticum*” CL12^T^, “*Peteryoungia ipomoeae*” shin9-1^T^, “*Peteryoungia rhizophila*” 7209-2^T^, “*Peteryoungia rosettiformans*” W3^T^, and “*Peteryoungia wuzhouensis*” W44^T^) formed a monophyletic group with 100% bootstrap support (**Figure 1**). All pairwise cpAAI values within this group were > 88%, while all pairwise cpAAI values against the other six “*R. aggregatum* complex” species were < 84.7% (**Figure 3**). These results support the proposal for the genus “*Peteryoungia*” [27], which should also primarily include *R. aggregatum*, as well as “*A. albertimagni*”, and “*R. glycinendophyticum*” once their names are validly published.

The remaining six “*R. aggregatum* complex” species formed a monophyletic group that could be further sub-divided into two clades. One clade corresponded to the three *Ciceribacter* species, while the other clade contained *R. daejeonense* DSM 17795^T^, *R. naphthalenivorans* TSY03b^T^, and *R. selenitireducens* ATCC BAA-1503^T^ (**Figure 1**). All within-group cpAAI values were > 86.5% while all between-group cpAAI values were ≤ 85.4%, providing support for these two clades representing separate genera. However, the bootstrap support for the split of these two clades in the phylogeny is only 80%, and the topology of the tree in this region (**Figure 1**) differs from the tree reported by Rahi *et al*., wherein *R. daejeonense, R. naphthalenivorans*, and *R. selenitireducens* were not monophyletic (see Figure 2 of [27]). Overall, the data presented here are not in agreement with the proposal to transfer *R. daejeonense, R. naphthalenivorans*, and *R. selenitireducens* to the genus *Ciceribacter*. Instead, we propose this clade be referred to as the “*R. daejeonense* complex” pending further study – enabled by the availability of additional genomes of strains belonging to these clades – to resolve whether these species belong to a novel genus or whether they should be transferred to the genus *Ciceribacter*.

### Proposal for the emendation of the genus *Sinorhizobium* as a distinct genus from *Ensifer*

Taxonomy of the *Ensifer* / *Sinorhizobium* genus has been the subject of discussion since the early 2000s. The genus *Ensifer* was proposed in 1982 to describe *Ensifer adhaerens*, a bacterial predator [8]. Subsequently, the genus *Sinorhizobium* was proposed in 1988 when *Rhizobium fredii* was reclassified as *Sinorhizobium fredii* [9], which was followed by the emendation of this genus by de Lajudie *et al*., 1994 [28]. In 2002, as the 16S rRNA gene sequence of *E. adhaerens* became available, the Subcommittee on the Taxonomy of *Agrobacterium* and *Rhizobium* (hereafter “the subcommittee”) of the International Committee on Systematics of Prokaryotes (ICSP) noted that this taxon is a part of *Sinorhizobium* [29]. Although the subcommittee pointed out that the name *Ensifer* has priority, conservation of the name *Sinorhizobium* was endorsed in contravention of the rules of the International Code of Nomenclature of Prokaryotes (ICNP). Neighbour-joining trees constructed from 16S rRNA gene sequences or partial *recA* gene sequences, together with phenotypic data, provided further data interpreted as supporting the synonymy and unification of the genera *Sinorhizobium* and *Ensifer*, leading Willems *et al*. (2003) to propose the new combination “*Sinorhizobium adhaerens*” [30]. Accordingly, in their Request for an Opinion to the Judicial Commission, Willems *et al*. officially proposed to conserve the name *Sinorhizobium* [30]. As the primary argument for conservation of the name *Sinorhizobium*, the authors indicated that the name *Ensifer* would cause misunderstanding and confusion in the scientific community. A few months later, in his Request for an Opinion to the Judicial Commission, J. M. Young argued that *Ensifer*, not *Sinorhizobium*, was the valid name for the unified genus, as *Ensifer* had priority [31]. At the same time, J. M. Young emended the description of the genus *Ensifer*, and transferred previously described *Sinorhizobium* species to this genus [31]. The Judicial Commission of the ICSP (Judicial Opinion 84) later confirmed that *Ensifer* had priority over *Sinorhizobium*, pointed out that the name “*Sinorhizobium adhaerens*” is not validly published, and supported the transfer of members of the genus *Sinorhizobium* to *Ensifer* [32]. In this Opinion, it was claimed that the transfer of the members of the genus *Sinorhizobium* to the genus *Ensifer* would not cause confusion. The subcommittee, however, disagreed with this justification [33]. J. M. Young criticized these actions of the subcommittee [34], which was also later acknowledged by Tindall [35]. As predicted by Willems *et al*. [30], adoption of the genus name *Ensifer* continues to be met with resistance from many rhizobiologists [36].

Earlier phylogenetic studies noted that *E. adhaerens* was an outgroup of the genus *Ensifer* [37], providing some support that *E. adhaerens* represented a distinct genus; however, it was suggested that further evidence would be required prior to redefining genera within this clade [37]. Significant phylogenomic and phenotypic data now exists providing strong evidence that the genera *Ensifer* and *Sinorhizobium* as defined Casida 1982 [8] and Chen *et al*. 1988 [9], respectively, refer to closely related, yet separate, taxa. At least seven studies, including the Genome Taxonomy Database, have presented phylogenetic trees containing two well-defined clades within the genus *Ensifer* [20, 25, 38–42]. These phylogenies were built on the basis of gene (up to 1,652 genes) or protein (up to 155 proteins) sequences using ML or Bayesian inference analysis approaches, indicating that the observed clades are robust to the choice of phylogenetic approach. Notably, our recent study presents a ML phylogeny where the genus *Ensifer* is split into two clades of 12 and 20 genospecies with 100% bootstrap support for the split, which we the defined as the “nonsymbiotic” and “symbiotic” clades, respectively [20]. The split is also observed in a ML phylogeny of the 16S rRNA genes of the same strains, with 62% bootstrap support (**Figure S8**). We similarly see a split of the genus *Ensifer* into two clades of three species type strains (including *E. adhaerens* Casida A^T^, the type strain of the type species of the genus *Ensifer* Casida 1982) and 12 species type strains (including *E. fredii* USDA 205^T^, the type strain of the type species of the genus *Sinorhizobium* Chen *et al*. 1988) in our core-genome gene phylogeny, with 100% bootstrap support (**Figure 1**), representing the nonsymbiotic and symbiotic clades, respectively. However, all between-clade cpAAI values were above the suggested 86% threshold as a baseline criterion for genus delimitation. Despite this, and following our proposed framework, we argue that there is sufficient other genomic and phenotypic data supporting the division of this genus (cf. Figures 3, 4, and 5 of [20]). We describe the distinctive traits and respective synapomorphies of these clades in **Table 1** and below.

**Figure S8.**
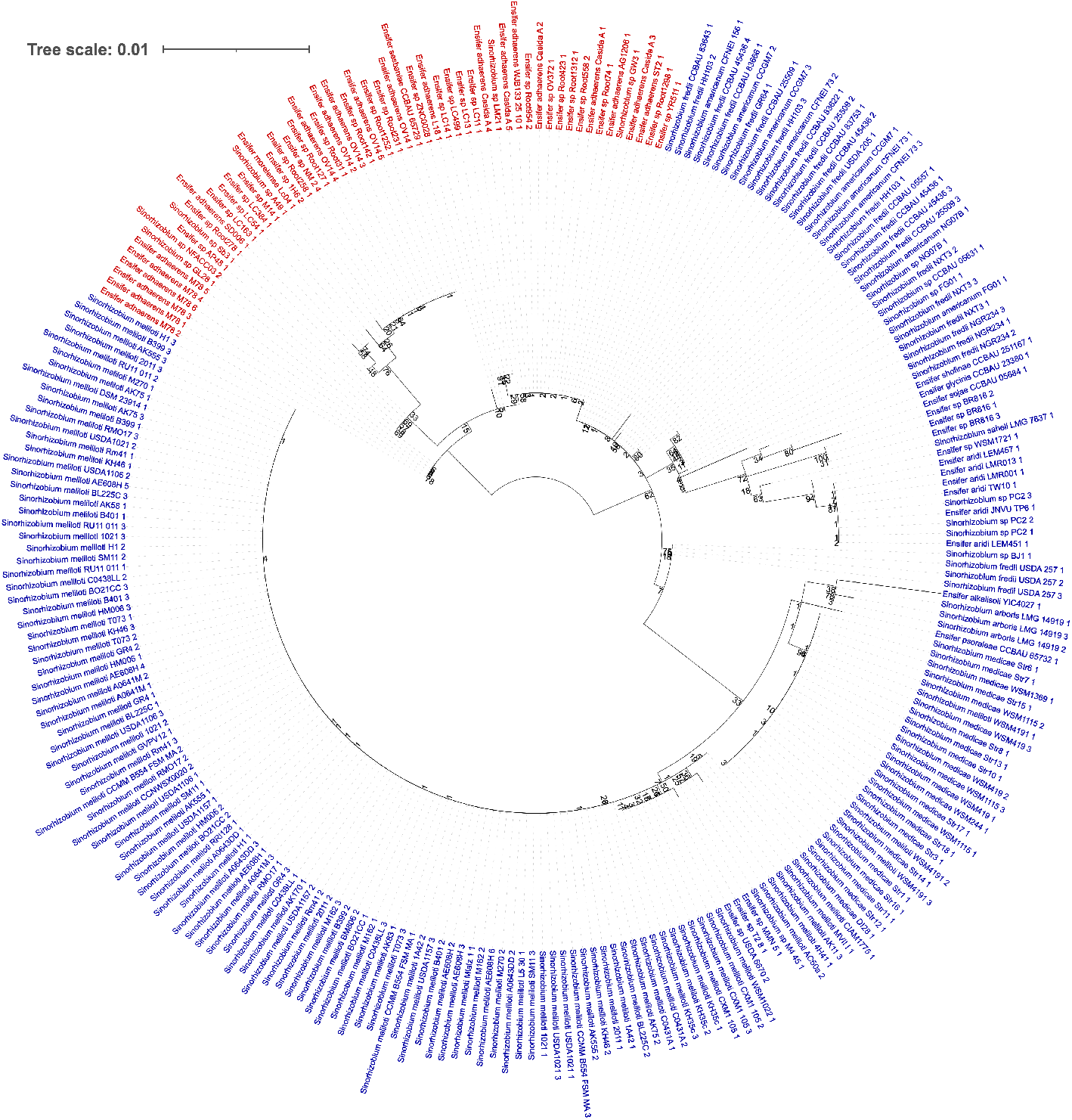
16S rRNA gene phylogeny of the genera *Ensifer* and *Sinorhizobium*. A maximum likelihood phylogeny of 258 16S rRNA genes from 157 strains of the genera *Ensifer* (red) and *Sinorhizobium* (blue) is shown. Strains are named as recorded in NCBI at the time of collection. The phylogeny was created using RAxML as described in the Materials and Methods. Values represent bootstrap support from 756 bootstrap replicates. The scale represents the mean number of nucleotide substitutions per site. One 16S rRNA gene from each of *E*. sp. GL28 and *S. meliloti* BM806 are not shown due to extremely long branch lengths.

**Table 1.**
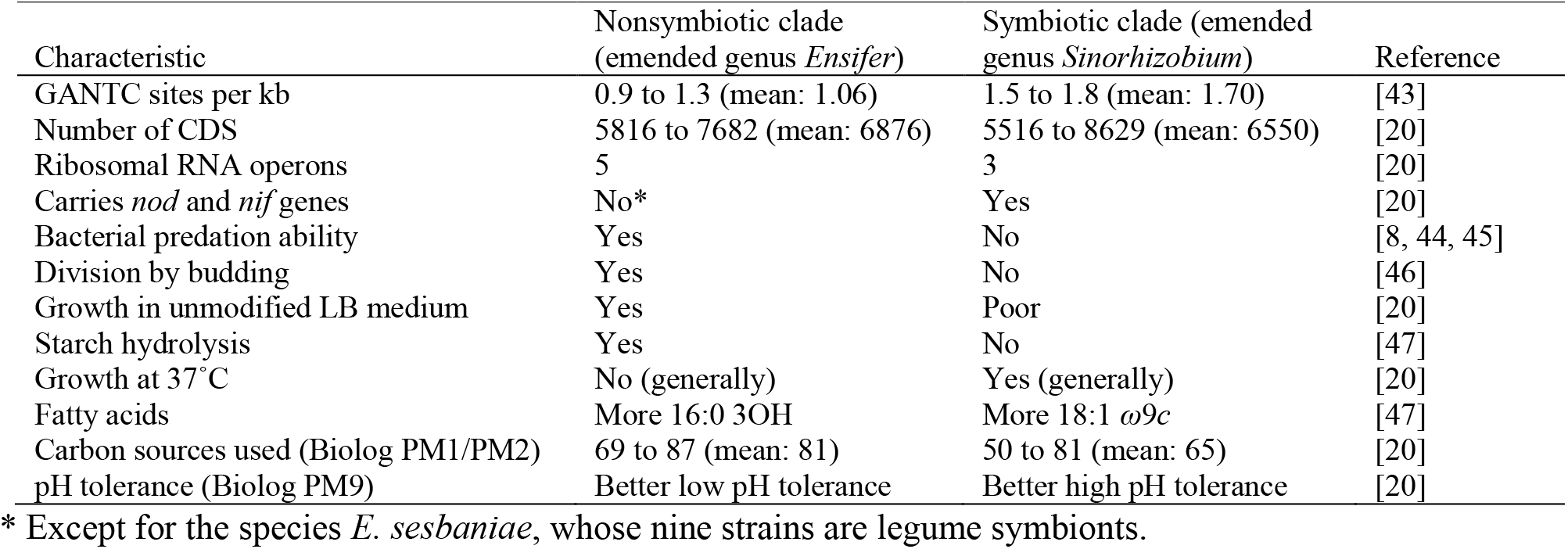
Characteristics differentiating the previously-defined nonsymbiotic and symbiotic clades of the genus *Ensifer*, corresponding to the emended genera *Ensifer* and *Sinorhizobium*, respectively.

The genome-wide frequency of the pentanucleotide GANTC is higher in all genomes of the symbiotic clade compared to all genomes of the nonsymbiotic clade, with a statistically significant average difference of 60% (1.70 vs 1.06 GANTC sites per kb, *p-*-value < 1 × 10^−10^ using a two-sample *t*-test) [43]. As the GANTC motif is methylated by the highly conserved cell cycle-regulated methyltransferase CcrM [48, 49], this difference may reflect an important difference in the cell cycle biology of these two clades [43]. Indeed, species of the nonsymbiotic clade (*E. adhaerens* and *E. morelensis*) are capable of division by budding unlike species of the symbiotic clade [46]. It has also been shown that the ability to hydrolyze starch [47] and robustly grow in LB broth lacking Ca^2+^ and Mg^2+^ ion supplementation [20] is specific to the nonsymbiotic clade. Stress tolerance of the two clades also differs (based on an analysis of 10 representative strains), with strains of the nonsymbiotic clade generally being more tolerant to alkaline conditions while strains of the symbiotic clade were generally more acid-tolerant and heat-tolerant [20]. Although many catabolic abilities could be found in at least a subset of each clade, which is unsurprising given both clades have open pangenomes, species of the nonsymbiotic clade are capable of catabolizing an average of 81 (out of 190 tested) carbon sources compared to an average of 65 for the symbiotic clade [20]. These differences in general phenotypic traits, together with the additional genomic and phenotypic differences outlined in **Table 1**, indicate marked differences in the biology of strains from these two clades. Indeed, at least two genospecies of the nonsymbiotic clade have been described as bacterial predators [8, 44, 45], a lifestyle that has not been attributed to any members of the symbiotic clade. Moreover, these two clades display significant differences in relation to their interactions with plant species, specifically, a biased distribution of the *nod* and *nif* genes required for establishment of nitrogen-fixing symbiosis with legumes [20, 41]. A recent study showed that whereas the core *nodABC* and *nifHDK* genes were present in strains from all 20 genospecies of the symbiotic clade, they were observed in just one of the 12 genospecies belonging the nonsymbiotic clade (*E. sesbaniae*, with all 9 reported strains, isolated from three different geographic origins, being symbiotic) [20, 47]. Symbiotic traits are linked to the presence of an accessory megaplasmid in the genome, and thus should not be considered relevant in delineating taxa [6]. However, this almost unique ability of genomes from the symbiotic clade to host symbiotic megaplasmids with respect to their relatives from the nonsymbiotic clade likely reflects differences in their genetic background. These discrepancies in symbiotic potential could thus be interpreted as a further marker of differentiated biology between these two clades.

Taken together, these genomic and phenotypic data suggest that the organisms in these two clades significantly differ in their biology and ecology, reminiscent of the stable ecotype model for bacterial species [50]. Notably, the type species of the genus *Ensifer* (*E. adhaerens*) is found within the nonsymbiotic clade, while the original type species of the genus *Sinorhizobium* (*S. fredii*) is found within the symbiotic clade. Given we established the taxonomic position of these type species-containing clades to be well separated, we argue that the proposal of Willems *et al*. (2003) [30] to unify the genera *Ensifer* Casida 1982 and *Sinorhizobium* Chen *et al*. 1988, and the Judicial Opinion 84 enacting the transfer of the members of the genus *Sinorhizobium* to the genus *Ensifer* [32], are no longer supported. Instead, we propose that *Ensifer* Casida 1982 and *Sinorhizobium* Chen *et al*. 1988 refer to closely related sister genera, of which *Ensifer* and *Sinorhizobium*, respectively, are the legitimate names in accordance with Rules 51a and 23a of the ICNP [51]. We note that the subcommittee has previously indicated support for this proposal [52], while also stating they are not in favour of creating subgenera for these taxa [36]. Formal genus and species emendations and circumscriptions are provided below.

### Taxonomy of the genus *Neorhizobium*

More study is required to resolve the taxonomic relationships between the “*Neorhizobium sensu stricto*” clade (**Figure 1**) – which includes *N. vignae, N. alkalisolii, N. hautlense, N. galegae* and *N. tomejilense*, as well as *Rhizobium petrolearium* – and related taxa. The core-genome gene phylogeny (**Figure 1**) and the cpAAI data (**Figure 3**) suggest that “*Neorhizobium lilium*” represents a new genus, as does the clade formed by *Neorhizobium* sp. NCHU2750, *Rhizobium smilacinae*, and *Rhizobium cellulosilyticum*. However, because bootstrap values provided only moderately good support for the topology of the tree in the extended *Neorhizobium* clade and the clades were not well-resolved by the cpAAI data, we defer the proposal of new genera until publication of further genomic evidence.

### Additional taxonomic implications of the proposed framework for genus delimitation

*R. petrolearium* DSM 26482^T^ formed a clade with “*Neorhizobium sensu stricto*” (**Figure 1**). Pairwise cpAAI values were all > 90% when *R. petrolearium* was compared against “*Neorhizobium sensu stricto*” species type strains (**Figure 3**). We therefore propose that *R. petrolearium* be transferred to the genus *Neorhizobium* (see below for formal description). *Rhizobium tarimense* CCTC AB 2011022^T^ formed a clade with the genus *Pseudorhizobium* (**Figure 1**). Pairwise cpAAI values were all > 88% when *R. tarimense* was compared against the four *Pseudorhizobium* species type strains, but < 85% when compared against all other species (**Figure 3**). We therefore propose that *R. tarimense* be transferred to the genus *Pseudorhizobium* (see below for formal description).

*Rhizobium arenae* MIM27^T^ formed a clade with the genus *Pararhizobium*. Pairwise cpAAI values were 87.4%, 87.4%, and 85.7% when *R. arenae* was compared against the three *Pararhizobium* species type strains, but < 84% when compared against all other species (**Figure 3**). We therefore propose that *R. arenae* be transferred to the genus *Pararhizobium* (see below for formal description).

*Rhizobium azooxidifex* DSM 100211^T^ formed a clade with “*Mycoplana subbaraonis*” JC85^T^ and *Mycoplana dimorpha* DSM 7138^T^ (**Figure 1**). Pairwise cpAAI values between these three species type strains were all > 89%, while cpAAI values against strains outside of this clade were all < 85% (**Figure 3**). We therefore propose that *R. azooxidifex* be transferred to the genus *Mycoplana* (see below for formal description).

*Rhizobium yantingense* CCTCC AB 2014007^T^ formed a clade with *Endobacterium cereale* (**Figure 1**). The pairwise cpAAI value between these two species was > 95%, while cpAAI values against strains outside of this clade were all < 84% (**Figure 3**). We therefore propose that *R. yantingense* be transferred to the genus *Endobacterium* (see below for formal description).

To confirm the distinct taxonomic positions of the above-mentioned species and support their transfer to the respective genera, we compared them to other genus members using ANIb and isDDH indices. These indices are regarded as standard measures of relatedness between prokaryotic species that were widely used for species delimitation [53, 54]. In all cases, the obtained values were clearly below the thresholds for species delimitation (95–96% for ANI or 70% for DDH) (**Datasets S3 and S4**), confirming the authenticity of these species.

In addition, the following species are candidates as type species for new genera: “*R. album*”, *R. populii, R. borbori*, and *R. halophytocola*. The reclassification of *R. album* into a new genus was also suggested by Young *et al*. 2021 [5]. Moreover, *R. helianthi* CGMC 1.12192^T^, *R. rhizoryzae* DSM 29514^R^, and *Allorhizobium pseudoryzae* DSM 19479^T^ formed a monophyletic group as a sister taxon to the genus *Endobacterium* (**Figure 1**). This clade of three species type strains is another candidate for reclassification as a new genus as pairwise cpAAI values within the clade were all > 86.5% while all cpAAI values with species outside of the clade were all < 83% (**Figure 3**).

### Description of *Xaviernesmea* gen. nov

*Xaviernesmea* (gza.vje.nem’e.a.; N.L. fem. n., in honour of Dr. Xavier Nesme, taxonomist of agrobacteria and rhizobia who pioneered the use of reverse ecology approaches to infer the ecology of *Agrobacterium* genomic species from comparative genomic analyses [7]).

Cells are Gram-negative, rod-shaped, and aerobic. Oxidase and catalase positive. Can utilise adonitol, D-raffinose and succinic acid and pH range for growth is 5.0–11.0 [55]. The G+C content of the genomic DNA is in the range 62.8-64.7 %. The genus *Xaviernesmea* has been separated from other *Rhizobiaceae* genera based on a core-genome phylogeny and whole- and core-proteome relatedness indexes (wpAAI and cpAAI).

The type species is *Xaviernesmea oryzae*.

### Description of *Xaviernesmea oryzae* comb. nov

*Xaviernesmea oryzae* (o.ry’zae. L. gen. fem. n. *oryzae*, of rice, referring to the host of isolation of the type strain).

Basonym: *Rhizobium oryzae* Peng *et al*. 2008 [56].

Homotypic synonym: *Allorhizobium oryzae* (Peng et al., 2008) Mousavi *et al*. 2015.

The description is as provided by Peng *et al*. 2008 [56] and Mousavi *et al*. 2015 [26]. *X. oryzae* can be differentiated from other species of the genus *Xaviernesmea* based on OGRI calculations (ANI and isDDH). The genomic G+C content of the type strain is 62.8 %. Its approximate genome size is 5.39 Mbp.

The type strain is Alt 505^T^ (= LMG 24253^T^ = CGMCC 1.7048^T^), isolated from *Oryza alta* growing in the Wild Rice Core Collection Nursery of South China Agricultural University. The NCBI RefSeq Assembly accession number for the genome sequence is GCF_001938935.1.

### Description of *Xaviernesmea rhizosphaerae* sp. nov

*Xaviernesmea rhizosphaerae* (rhi.zo.sphae’rae. N.L. gen. fem. n. *rhizosphaerae*, of the rhizosphere, referring to host plant compartment of isolation of the type strain).

The description is as provided by Zhao *et al*. 2017 [55]. *X. rhizosphaerae* can be differentiated from other species of the genus *Xaviernesmea* based on OGRI calculations (ANI and isDDH). The genomic G+C content of the type strain is 64.7 %. Its approximate genome size is 5.18 Mbp.

The type strain is MH17^T^ (= ACCC 19963^T^ = KCTC 52414^T^), which was isolated from root of rice collected from Beijing, China. The NCBI RefSeq Assembly accession number for the genome sequence is GCF_001938945.1. We note that the name “*Rhizobium rhizosphaerae*” that was proposed in the original publication [55] has yet to be validly published.

### Emended description of the genus *Ensifer* Casida 1982

The description is as given by Casida 1982 [8] with the following emendations. The optimal growth temperature is 27-28°C. Some species can grow at 37°C. Capable of growth in unmodified Lysogeny Broth (LB). Capable of hydrolyzing starch. Resistant to multiple antibiotics including ampicillin and erythromycin. The genomic G+C content is ∼ 61-63 %. The genomic GANTC pentanucleotide frequency is ∼ 0.9-1.3 sites per kb. Most strains carry 5 rRNA operons. The genus can be differentiated from other genera based on core-genome gene phylogenies.

The type species is *Ensifer adhaerens*.

The emended genus contains the species *E. adhaerens, E. morelensis*, and *E. sesbaniae*.

The species *E. alkalisoli, E. americanum, E. arboris, E. fredii, E. garamanticum, E. glycinis E. kostiensis, E. kummerowiae, E. medicae, E. meliloti, E. mexicanum, E. numidicum, E. psoraleae, E. saheli, E. sofinae, E. sojae*, and *E. terangae* are transferred to the genus *Sinorhizobium*.

### Emended description of the genus *Sinorhizobium* Chen et al. 1988 emend. de Lajudie *et al*. 1994

The description is as given by de Lajudie *et al*. 1994 [28] with the following emendations, drawing also from Young 2003 [31]. The optimal growth temperature is 25-33°C, but some strains can grow at 12°C and others can grow at 44°C. Optimum pH is 6-7, but some strains can grow at pH 5.0 and others at pH 10.0. Starch is not utilized. Ammonium salts, nitrate, nitrite, and many amino acids can serve as nitrogen sources for most strains. Most strains produce cytochrome oxidase and catalase. The genomic G+C content is ∼ 61-64 %. The genomic GANTC pentanucleotide frequency is ∼ 1.5-1.8 sites per kb. Most strains carry 3 rRNA operons. The genus can be differentiated from other genera based on core-genome gene phylogenies.

The type species is *Sinorhizobium fredii*.

The emended genus contains the species *S. alkalisoli, S. americanum, S. arboris, S. fredii, S. garamanticum, S. glycinis, S. kostiense, S. kummerowiae, S. medicae, S. meliloti, S. mexicanum, S. numidicum, S. psoraleae, S. saheli, S. sofinae, S. sojae*, and *S. terangae*.

### Description of *Sinorhizobium alkalisoli* comb. nov

*Sinorhizobium alkalisoli* (al.ka.li.so’li. N.L. neut. n. *alkali*, alkali (from Arabic al-qaliy); L. neut. n. *solum*, soil; N.L. gen. neut. n. *alkalisoli*, of alkaline soil, referring to the saline-alkali soil where the bacterium was isolated).

Basonym: *Ensifer alkalisoli* Li *et al*. 2016.

The description is as provided by Li *et al*. 2016 [57]. *S. alkalisoli* can be differentiated from other species of the genus *Sinorhizobium* based on OGRI calculations (ANI and isDDH). The genomic G+C content of the type strain is 62.2 %. Its approximate genome size is 6.13 Mbp.

The type strain is YIC4027^T^ (=HAMBI 3655^T^ =LMG 29286^T^). The NCBI RefSeq Assembly accession number for the genome sequence is GCF_008932245.1.

### Emended description of *Sinorhizobium americanum* Toledo *et al*. 2004

*Sinorhizobium americanum* (a.me.ri.ca’num. N.L. neut. adj. *americanum*, American, referring to the isolation of the type strain from the Colorado Plateau).

Homotypic synonym: *Ensifer americanus* (Toledo *et al*. 2004) Wang et al. 2015 emend. Hördt et al. 2020.

The description is as provided by Hördt et al. 2020 [4]. *S. americanum* can be differentiated from other species of the genus *Sinorhizobium* based on OGRI calculations (ANI and isDDH). The genomic G+C content of the type strain is 62.3 %. Its approximate genome size is 6.75 Mbp.

The type strain is CFNEI 156^T^ (=ATCC BAA-532^T^ =CIP 108390^T^ =DSM 15007^T^). The NCBI RefSeq Assembly accession number for the genome sequence is GCF_001889105.1.

### Emended description of *Sinorhizobium arboris* Nick *et al*. 1999

*Sinorhizobium arboris* (ar’bo.ris. L. fem. n. *arbor*, tree; L. gen. fem. n. *arboris*, of a tree).

Homotypic synonym: *Ensifer arboris* (Nick *et al*. 1999) Young 2003 emend. Hördt et al. 2020.

The description is as provided by Hördt et al. 2020 [4]. *S. arboris* can be differentiated from other species of the genus *Sinorhizobium* based on OGRI calculations (ANI and isDDH). The genomic G+C content of the type strain is 62.0 %. Its approximate genome size is 6.85 Mbp.

The type strain is HAMBI 1552^T^ (=ATCC BAA-226^T^ =DSM 13375^T^ =LMG 14919^T^ =NBRC 100383^T^ =TTR 38^T^). The NCBI RefSeq Assembly accession number for the genome sequence is GCF_000427465.1.

### Emended description of *Sinorhizobium fredii* (Scholla and Elkan 1984) Chen *et al*. 1988

*Sinorhizobium fredii* (fred’i.i. N.L. gen. neut. n. *fredii*, of Fred, named after of E.B. Fred).

Homotypic synonym: *Ensifer fredii* (Scholla and Elkan 1984) Young 2003 emend. Hördt et al. 2020.

Heterotypic synonym: *Ensifer xinjiangensis* (Chen *et al*. 1988) Young 2003 [39]

The description is as provided by Hördt et al. 2020 [4]. *S. fredii* can be differentiated from other species of the genus *Sinorhizobium* based on OGRI calculations (ANI and isDDH). The genomic G+C content of the type strain is 62.2 %. Its approximate genome size is 7.15 Mbp.

The type strain is USDA 205^T^ (=ATCC 35423^T^ =CCUG 27877^T^ =DSM 5851^T^ =HAMBI 2075^T^ =ICMP 11139^T^ =IFO 14780^T^ =JCM 20967^T^ =LMG 6217^T^ =NBRC 14780^T^ =NRRL B-14241^T^ =NRRL B-14594^T^ =PRC 205^T^). The NCBI RefSeq Assembly accession number for the genome sequence is GCF_009601405.1.

### Description of *Sinorhizobium garamanticum* comb. nov

*Sinorhizobium garamanticum* (ga.ra.man’ti.cum. N.L. neut. adj. *garamanticum*, pertaining to Garamante, Garamantian, the country of Garamantes, from which the strains were isolated).

Basonym: *Ensifer garamanticus* Merabet *et al*. 2010.

The description is as provided by Merabet *et al*. 2010 [58]. *S. garamanticum* can be differentiated from other species of the genus *Sinorhizobium* by phylogenetic analysis based on several housekeeping (*recA, glnA, gltA, thrC* and *atpD*) genes and 16S rRNA gene sequencing. The genomic G+C content of the type strain is approximately 62.4 % (HPLC).

The type strain is ORS 1400^T^ (=CIP 109916^T^ =LMG 24692^T^).

### Description of *Sinorhizobium glycinis* comb. nov

*Sinorhizobium glycinis* (gly.ci’nis. N.L. gen. n. *glycinis*, of the botanical genus *Glycine*, the soybean, named for its nodulation characteristics and symbiotic genes).

Basonym: *Ensifer glycinis* Yan *et al*. 2016.

The description is as provided by Yan *et al*. 2016 [59]. *S. glycinis* can be differentiated from other species of the genus *Sinorhizobium* based on OGRI calculations (ANI and isDDH). The genomic G+C content of the type strain is 62.4 %. Its approximate genome size is 6.04 Mbp.

The type strain is CCBAU 23380^T^ (=HAMBI 3645^T^ =LMG 29231^T^). The NCBI RefSeq Assembly accession number for the genome sequence is GCF_001651865.1.

### Emended description of *Sinorhizobium kostiense* Nick *et al*. 1999

*Sinorhizobium kostiense* (kos.ti.en’se. N.L. masc./fem. adj. *kostiense*, pertaining to Kosti, the region in Sudan where most of these organisms have been isolated).

Homotypic synonym: *Ensifer kostiensis* (Nick *et al*. 1999) Young 2003.

The description is as provided by Nick *et al*. 1999 [60]. *S. kostiense* can be differentiated from other species of the genus *Sinorhizobium* based on OGRI calculations (ANI and isDDH). The genomic G+C content of the type strain is 61.7 %. Its approximate genome size is 6.33 Mbp.

The type strain is DSM 13372^T^ (=ATCC BAA-227^T^ =HAMBI 1489^T^ =LMG 15613^T^ =LMG 19227^T^ =NBRC 100382^T^ =TTR 15^T^). The NCBI RefSeq Assembly accession number for the genome sequence is GCF_001651865.1.

### Emended description of *Sinorhizobium kummerowiae* Wei *et al*. 2002

*Sinorhizobium kummerowiae* (kum.me.ro’wi.ae. N.L. gen. fem. n. *kummerowiae*, of *Kummerowia*, a genus of leguminous plants, referring to the host from which the bacterium was isolated).

Homotypic synonym: *Ensifer kummerowiae* (Wei *et al*. 2002) Young 2003.

The description is as provided by Wei *et al*. 2002 [61]. *S. kummerowiae* can be differentiated from other species of the genus *Sinorhizobium* by phylogenetic analysis based on 16S rRNA gene sequences. The genomic G+C content of the type strain is approximately 61.6 % (*Tm*).

The type strain is CCBAU 71714^T^ (=CGMCC 1.3046^T^ =CIP 108026 ^T^ =NBRC 100799^T^).

### Emended description of *Sinorhizobium medicae* Rome *et al*. 1996

*Sinorhizobium medicae* (me’di.cae. L. gen. fem. n. *medicae*, of/from lucern (plant belonging to the genus *Medicago*).

Homotypic synonym: *Ensifer medicae* (Rome et al. 1996) Young 2003.

The description is as provided by Rome *et al*. 1996 [62]. *S. medicae* can be differentiated from other species of the genus *Sinorhizobium* based on OGRI calculations (ANI and isDDH). The genomic G+C content of the type strain is 61.2 %. Its approximate genome size is 6.53 Mbp.

The type strain is A 321^T^ (=11-3 21a^T^ =HAMBI 2306^T^ =ICMP 13798^T^ =LMG 19920^T^ =NBRC 100384^T^ =USDA 1037^T^). The NCBI RefSeq Assembly accession number for the genome sequence is GCF_007827695.

### Emended description of *Sinorhizobium meliloti* (Dangeard 1926) de Lajudie *et al*. 1994

*Sinorhizobium meliloti* (me.li.lo’ti. N.L. masc. n. *Melilotus*, generic name of sweet clover; N.L. gen. masc. n. *meliloti*, of *Melilotus*).

Homotypic synonym: *Ensifer meliloti* (Dangeard 1926) Young 2003.

The description is as provided by de Lajudie *et al*. 1994 [28]. *S. meliloti* can be differentiated from other species of the genus *Sinorhizobium* based on OGRI calculations (ANI and isDDH). The genomic G+C content of the type strain is 62.0 %. Its approximate genome size is 7.34 Mbp.

The type strain is USDA 1002^T^ (=ATCC 9930^T^ =CCUG 27879^T^ =CFBP 5561^T^ =DSM 30135^T^ =HAMBI 2148^T^ =ICMP 12623^T^ =IFO 14782^T^ =JCM 20682^T^ =LMG 6133^T^ =NBRC 14782 =NCAIM B.01520^T^ =NRRL L-45^T^ =NZP 4027^T^). The NCBI RefSeq Assembly accession number for the genome sequence is GCF_007827695.1.

### Description of *Sinorhizobium mexicanum* comb. nov

*Sinorhizobium mexicanum* (me.xi.ca’num. N.L. neut. adj. *mexicanum*, of or belonging to Mexico, where the strains were isolated).

Basonym: *Ensifer mexicanus* Lloret *et al*. 2011.

The description is as provided by Lloret *et al*. 2011 [63]. *S. mexicanum* can be differentiated from other species of the genus *Sinorhizobium* based on OGRI calculations (ANI and isDDH). The genomic G+C content of the type strain is 61.4 %. Its approximate genome size is 7.14 Mbp.

The type strain is ITTG R7^T^ (=ATCC BAA-1312^T^ =CFN ER1001^T^ =CIP 109033^T^ =DSM 18446^T^ =HAMBI 2910^T^). The NCBI Assembly accession number for the genome sequence is GCF_013488225.1.

### Description of *Sinorhizobium numidicum* comb. nov

*Sinorhizobium numidicum* (nu.mi’di.cum. N.L. neut. adj. *numidicum*, pertaining to the country of Numidia, Numidian, the Roman denomination of the region in North Africa from which the majority of the organisms were isolated).

Basonym: *Ensifer numidicus* Merabet *et al*. 2010.

The description is as provided by Merabet *et al*. 2010 [58]. *S. numidicus* can be differentiated from other species of the genus *Sinorhizobium* by phylogenetic analysis based on several housekeeping (*recA, glnA, gltA, thrC* and *atpD*) genes and 16S rRNA gene sequencing. The genomic G+C content of the type strain is approximately 62.8 % (HPLC).

The type strain is ORS 1407^T^ (=CIP 109916^T^ =LMG 24692^T^).

### Description of *Sinorhizobium psoraleae* comb. nov

*Sinorhizobium psoraleae* (pso.ra.le’a.e N.L. gen. fem. n. *psoraleae*, of *Psoralea*, referring to the main host of the species).

Basonym: *Ensifer psoraleae* Wang *et al*. 2013.

The description is as provided by Wang *et al*. 2013 [47]. *S. psoraleae* can be differentiated from other species of the genus *Sinorhizobium* based on OGRI calculations (ANI and isDDH). The genomic G+C content of the type strain is 61.3 %. Its approximate genome size is 7.43 Mbp.

The type strain is CCBAU 65732^T^ (=HAMBI 3286^T^ =LMG 26835^T^). The NCBI RefSeq Assembly accession number for the genome sequence is GCF_013283645.1.

### Emended description of *Sinorhizobium saheli* de Lajudie *et al*. 1994

*Sinorhizobium saheli* (sa’hel.i. N.L. gen. neut. n. *saheli*, of the Sahel, the region in Africa from which they were isolated).

Homotypic synonym: *Ensifer saheli* (de Lajudie *et al*. 1994) Young 2003 emend. Hördt et al. 2020.

The description is as provided by Hördt et al. 2020 [4]. *S. saheli* can be differentiated from other species of the genus *Sinorhizobium* based on OGRI calculations (ANI and isDDH). The genomic G+C content of the type strain is 63.6 %. Its approximate genome size is 5.99 Mbp.

The type strain is LMG 7837^T^ (=ATCC 51690^T^ =DSM 11273^T^ =HAMBI 215^T^ =ICMP 13648^T^=NBRC 100386^T^ =ORS 609^T^). The NCBI RefSeq Assembly accession number for the genome sequence is GCF_001651875.1.

### Description of *Sinorhizobium shofinae* comb. nov

*Sinorhizobium shofinae* (sho.fi’nae. N.L. fem. gen. n. *shofinae* from Shofine, a company name, referring to the fact that the type strain of this species was isolated from root nodule of soybean grown in the farm of the company, Shandong Shofine Seed Technology Company Ltd., located in Jiaxiang County, Shandong Province of China).

Basonym: *Ensifer shofinae* Chen *et al*. 2017 emend. Hördt et al. 2020.

The description is as provided by Hördt et al. 2020 [4]. *S. shofinae* can be differentiated from other species of the genus *Sinorhizobium* based on OGRI calculations (ANI and isDDH). The genomic G+C content of the type strain is 59.9 %. Its approximate genome size is 6.21 Mbp.

The type strain is CCBAU 251167^T^ (=HAMBI 3507^T^ =LMG 29645^T^ =ACCC 19939^T^). The NCBI RefSeq Assembly accession number for the genome sequence is GCF_001704765.1.

### Description of *Sinorhizobium sojae* comb. nov

*Sinorhizobium sojae* (so’ja.e. N.L. gen. n. sojae, of soja, of soybean, referring to the source of the first isolates).

Basonym: *Ensifer sojae* Li *et al*. 2011 emend. Hördt et al. 2020.

The description is as provided by Hördt et al. 2020 [4]. *S. sojae* can be differentiated from other species of the genus *Sinorhizobium* based on OGRI calculations (ANI and isDDH). The genomic G+C content of the type strain is 60.9 %. Its approximate genome size is 6.09 Mbp.

The type strain is CCBAU 5684^T^ (=DSM 26426^T^ =HAMBI 3098^T^ =LMG 25493^T^). The NCBI RefSeq Assembly accession number for the genome sequence is GCF_002288525.1.

### Emended description of *Sinorhizobium terangae* de Lajudie et al. 1994

*Sinorhizobium terangae* (te’ran.gae. N.L. n. *terengae*, hospitality (from West African Wolof n. terenga, hospitality); N.L. gen. n. *terangae*, of hospitality, intended to mean that this species is isolated from different host plants).

Homotypic synonym: *Ensifer terangae* (de Lajudie et al. 1994) Young 2003.

The description is as provided by de Lajudie *et al*. 1994 [28]. *S. terangae* can be differentiated from other species of the genus *Sinorhizobium* based on OGRI calculations (ANI and isDDH). The genomic G+C content of the type strain is 61.4 %. Its approximate genome size is 6.79 Mbp.

The type strain is SEMIA 6460^T^ (=ATCC 51692^T^ =DSM 11282^T^ =HAMBI 220^T^ =ICM 13649^T^ =LMG 7834^T^ =NBRC 100385^T^ =ORS 1009^T^). The NCBI RefSeq Assembly accession number for the genome sequence is GCF_014197705.1.

### Description of *Endobacterium yantingense* comb. nov

*Endobacterium yantingense* (yan. ting. en’se. N.L. neut. adj. yantingense referring to Yanting district, Sichuan Province, PR China, where the organism was isolated).

Basonym: *Rhizobium yantingense* Chen *et al*. 2015.

The description is as provided by Chen *et al*. 2015 [64]. *E. yantingense* can be differentiated from another species of the genus *Endobacterium* (*Endobacterium cereale* corrig. Menéndez *et al*. 2021) based on OGRI calculations (ANI and isDDH). The genomic G+C content of the type strain is 59.5 %. Its approximate genome size is 5.82 Mbp.

The type strain is H66^T^ (=CCTCC AB 2014007^T^ =LMG 28229^T^), which was isolated from the surfaces of weathered rock (purple siltstone) in Yanting (Sichuan, PR China). The JGI IMG accession number for the genome sequence is Ga0196656.

### Description of *Neorhizobium petrolearium* comb. nov

*Neorhizobium petrolearium* (pe.tro.le.a’ri.um. L. fem. n. *petra*, rock; L. neut. *olearium* related to oil, of or belonging to oil; N.L. neut. adj. *petrolearium* related to mineral oil).

Basonym: *Rhizobium petrolearium* Zhang *et al*. 2012.

The description is as provided by Zhang *et al*. 2012 [65]. *N. petrolearium* can be differentiated from another species of the genus *Neorhizobium* based on OGRI calculations (ANI and isDDH). The genomic G+C content of the type strain is 60.5 %. Its approximate genome size is 6.97 Mbp. The type strain, SL-1^T^ (=ACCC 11238^T^ =KCTC 23288^T^), was isolated from petroleum-contaminated sludge samples in Shengli oilfield, Shandong Province, China. The JGI IMG accession number for the genome sequence is Ga0196653.

### Description of *Pararhizobium arenae* comb. nov

*Pararhizobium arenae* (a.re’nae. L. fem. gen. n. *arenae* of sand, the isolation source of the type strain).

Basonym: *Rhizobium arenae* Zhang *et al*. 2017.

The description is as provided by Zhang *et al*. 2017 [66]. *P. arenae* can be differentiated from another species of the genus *Pararhizobium* based on OGRI calculations (ANI and isDDH). The genomic G+C content of the type strain is 59.8 %. Its approximate genome size is 4.94 Mbp.

The type strain is MIM27^T^ (=KCTC 52299^T^ =MCCC 1K03215^T^), isolated from sand of the Mu Us Desert, PR China. The NCBI RefSeq Assembly accession number for the genome sequence is GCF_001931685.1.

### Description of *Pseudorhizobium tarimense* comb. nov

*Pseudorhizobium tarimense* (ta.rim.en’se. N.L. neut. adj. *tarimense*, pertaining to Tarim basin in Xinjiang Uyghur autonomous region of China, where the type strain was isolated).

Basonym: *Rhizobium tarimense* Turdahon *et al*. 2013.

The description is as provided by Turdahon *et al*. 2013 [67]. *P. tarimense* can be differentiated from other species of the genus *Pseudorhizobium* based on OGRI calculations (ANI and isDDH).

The genomic G+C content of the type strain is 61.2 %. Its approximate genome size is 4.83 Mbp.

The type strain is CCTCC AB 2011011^T^ (=NRRL B-59556^T^ =PL-41^T^). The JGI IMG accession number for the genome sequence is Ga0196649.

### Description of *Mycoplana azooxidifex* comb. nov

*Mycoplana azooxidifex* (a.zo.o.xi’di.fex. N.L. azooxidum, dinitrogenmonoxide; L. masc. suff. - fex, the maker; N.L. masc. n. azooxidifex, the dinitrogenmonoxide maker (nominative in apposition)).

Basonym: *Rhizobium azooxidifex* Turdahon *et al*. 2013.

The description is as provided by Turdahon *et al*. 2013 [67]. *M. azooxidifex* can be differentiated from other species of the genus *Mycoplana* based on OGRI calculations (ANI and isDDH). The genomic G+C content of the type strain is 64.3 %. Its approximate genome size is 5.89 Mbp.

The type strain is DSM 100211^T^ (=Po 20/26^T^ =LMG 28788^T^). The NCBI RefSeq Assembly accession number for the genome sequence is GCF_014196765.1.

## ACKNOWLEDGEMENTS

The authors gratefully acknowledge Prof. George M. Garrity (Michigan State University, East Lansing, MI, USA) for valuable help interpreting the ICNP, and Prof. Peter Young for helpful feedback on the manuscript. This research was enabled, in part, through computational resources provided by Compute Ontario (computeontario.ca), Compute Canada (computecanada.ca), and BMBF-funded de.NBI Cloud within the German Network for Bioinformatics Infrastructure (de.NBI) (031A537B, 031A533A, 031A538A, 031A533B, 031A535A, 031A537C, 031A534A, 031A532B). The work of NK was funded by the Deutsche Forschungsgemeinschaft (DFG, German Research Foundation) – Projektnummer 429677233. Research in the G.C.D laboratory is supported by a NSERC Discovery Grant, and funding from Queen’s University. A.M. and C.F. are supported by MICRO4Legumes grant (Italian Ministry of Agriculture) and ALL-IN project (H2020 ERA-NETs SUSFOOD2 and CORE Organic Cofund, under the Joint SUSFOOD2/CORE Organic Call 2019). This research was funded in part by the Wellcome Trust Grant [206194]. For the purpose of Open Access, the author has applied a CC-BY public copyright licence to any Author Accepted Manuscript version arising from this submission.

## DATASETS

**Dataset S1.** A list of strains included in our analyses, as well as the corresponding genome assembly accessions.

**Dataset S2.** A representative sequence for each of the 170 nonrecombinant genes used to construct the maximum likelihood phylogeny.

**Dataset S3.** Average nucleotide identity (ANIb) values for select taxa from the family *Rhizobiaceae*.

**Dataset S4.** *In silico* DNA-DNA hybridization (isDDH) values for select taxa from the family *Rhizobiaceae*.

## Notes

### Competing Interest Statement

The authors have declared no competing interest.

## REFERENCES

1. Conn HJ. Taxonomic relationships of certain non-sporeforming rods in soil. J Bacteriol 1938;36:320–321.

2. Alves LMC, Souza JAM de, Varani A de M, Lemos EG de M. The Family Rhizobiaceae. The Prokaryotes 2014;419–437.

3. Parte AC, Sardà Carbasse J, Meier-Kolthoff JP, Reimer LC, Göker M 2020. List of Prokaryotic names with Standing in Nomenclature (LPSN) moves to the DSMZ. Int J Syst Evol Microbiol;70:5607–5612.

4. Hördt A, López MG, Meier-Kolthoff JP, Schleuning M, Weinhold L-M, et al. Analysis of 1,000+ type-strain genomes substantially improves taxonomic classification of Alphaproteobacteria. Front Microbiol 2020;11:468.

5. Young JPW, Moeskjær S, Afonin A, Rahi P, Maluk M, et al. Defining the Rhizobium leguminosarum species complex. Genes 2021;12:111.

6. de Lajudie PM, Andrews M, Ardley J, Eardly B, Jumas-Bilak E, et al. Minimal standards for the description of new genera and species of rhizobia and agrobacteria. Int J Syst Evol Microbiol 2019;69:1852–1863.

7. Lassalle F, Muller D, Nesme X. Ecological speciation in bacteria: reverse ecology approaches reveal the adaptive part of bacterial cladogenesis. Res Microbiol 2015;166:729–741.

8. Casida LE. Ensifer adhaerens gen. nov., sp. nov.: a bacterial predator of bacteria in soil. Int J Syst Evol Microbiol 1982;32:339–345.

9. Chen WX, Yan GH, Li JL. Numerical taxonomic study of fast-growing soybean rhizobia and a proposal that Rhizobium fredii be assigned to Sinorhizobium gen. nov. Int J Syst Evol Microbiol 1988;38:392–397.

10. Contreras-Moreira B, Vinuesa P. GET_HOMOLOGUES, a versatile software package for scalable and robust microbial pangenome analysis. Appl Environ Microbiol 2013;79:7696–7701.

11. Vinuesa P, Ochoa-Sánchez LE, Contreras-Moreira B. GET_PHYLOMARKERS, a software package to select optimal orthologous clusters for phylogenomics and inferring pan-genome phylogenies, used for a critical geno-taxonomic revision of the genus Stenotrophomonas. Front Microbiol 2018;9:771.

12. Kuzmanović N, Biondi E, Overmann J, Puławska J, Verbarg S, et al. Revisiting the taxonomy of Allorhizobium vitis (i.e. Agrobacterium vitis) using genomics - emended description of All. vitis sensu stricto and description of Allorhizobium ampelinum sp. nov. bioRxiv. Epub ahead of print 21 December 2020. DOI: 10.1101/2020.12.19.423612.

13. Nguyen L-T, Schmidt HA, von Haeseler A, Minh BQ. IQ-TREE: a fast and effective stochastic algorithm for estimating maximum-likelihood phylogenies. Mol Biol Evol 2015;32:268–274.

14. Buchfink B, Xie C, Huson DH. Fast and sensitive protein alignment using DIAMOND. Nat Methods 2015;12:59–60.

15. Paradis E, Schliep K. ape 5.0: an environment for modern phylogenetics and evolutionary analyses in R. Bioinformatics 2019;35:526–528.

16. Qin Q-L, Xie B-B, Zhang X-Y, Chen X-L, Zhou B-C, et al. A proposed genus boundary for the prokaryotes based on genomic insights. J Bacteriol 2014;196:2210–2215.

17. Hyatt D, Chen G-L, Locascio PF, Land ML, Larimer FW, et al. Prodigal: prokaryotic gene recognition and translation initiation site identification. BMC Bioinform 2010;11:119.

18. Camacho C, Coulouris G, Avagyan V, Ma N, Papadopoulos J, et al. BLAST+: architecture and applications. BMC Bioinform 2009;10:421.

19. Meier-Kolthoff JP, Auch AF, Klenk H-P, Göker M. Genome sequence-based species delimitation with confidence intervals and improved distance functions. BMC Bioinform 2013;14:60.

20. Fagorzi C, Ilie A, Decorosi F, Cangioli L, Viti C, et al. Symbiotic and nonsymbiotic members of the genus Ensifer (syn. Sinorhizobium) are separated into two clades based on comparative genomics and high-throughput phenotyping. Genome Biol Evol 2020;12:2521–2534.

21. Katoh K, Standley DM. MAFFT multiple sequence alignment software version 7: improvements in performance and usability. Mol Biol Evol 2013;30:772–780.

22. Capella-Gutiérrez S, Silla-Martínez JM, Gabaldón T. trimAl: a tool for automated alignment trimming in large-scale phylogenetic analyses. Bioinformatics 2009;25:1972–1973.

23. Stamatakis A. RAxML version 8: a tool for phylogenetic analysis and post-analysis of large phylogenies. Bioinformatics 2014;30:1312–1313.

24. Zheng J, Wittouck S, Salvetti E, Franz CMAP, Harris HMB, et al. A taxonomic note on the genus Lactobacillus: Description of 23 novel genera, emended description of the genus Lactobacillus Beijerinck 1901, and union of Lactobacillaceae and Leuconostocaceae. Int J Syst Evol Microbiol 2020;70:2782–2858.

25. Lassalle F, Dastgheib SMM, Zhao F-J, Zhang J, Verbarg S, et al. Phylogenomics reveals the basis of adaptation of Pseudorhizobium species to extreme environments and supports a taxonomic revision of the genus. Syst Appl Microbiol 2021;44:126165.

26. Mousavi SA, Willems A, Nesme X, de Lajudie P, Lindström K. Revised phylogeny of Rhizobiaceae: Proposal of the delineation of Pararhizobium gen. nov., and 13 new species combinations. Syst Appl Microbiol 2015;38:84–90.

27. Rahi P, Khairnar M, Hagir A, Narayan A, Jain KR, et al. Peteryoungia gen. nov. with four new species combinations and description of Peteryoungia desertarenae sp. nov., and taxonomic revision of the genus Ciceribacter based on phylogenomics of Rhizobiaceae. Arch Microbiol. Epub ahead of print 9 May 2021. DOI: 10.1007/s00203-021-02349-9.

28. de Lajudie, Philippe P, Willems A, Pot B, Dewettinck D, Maestrojuan G, et al. Polyphasic taxonomy of rhizobia: emendation of the genus Sinorhizobium and description of Sinorhizobium meliloti comb. nov., Sinorhizobium saheli sp. nov., and Sinorhizobium teranga sp. nov. Int J Syst Evol Microbiol 1994;44:715–733.

29. Lindstrom K, Martínez-Romero ME. International Committee on Systematics of Prokaryotes: Subcommittee on the taxonomy of Agrobacterium and Rhizobium. Minutes of the meeting, 4 July 2001, Hamilton, Canada. Int J Syst Evol Microbiol 2002;52:2337.

30. Willems A, Fernández-López M, Muñoz-Adelantado E, Goris J, De Vos P, et al. Description of new Ensifer strains from nodules and proposal to transfer Ensifer adhaerens Casida 1982 to Sinorhizobium as Sinorhizobium adhaerens comb. nov. Request for an opinion. Int J Syst Evol Microbiol 2003;43:1207–1217.

31. Young JM. The genus name Ensifer Casida 1982 takes priority over Sinorhizobium Chen et al. 1988, and Sinorhizobium morelense Wang et al. 2002 is a later synonym of Ensifer adhaerens Casida 1982. Is the combination ‘Sinorhizobium adhaerens’ (Casida 1982) Willems et al. 2003 legitimate? Request for an opinion. Int J Syst Evol Microbiol 2003;53:2107–2110.

32. Judicial Commission of the International Committee on Systematics of Prokaryotes. The genus name Sinorhizobium Chen et al. 1988 is a later synonym of Ensifer Casida 1982 and is not conserved over the latter genus name, and the species name ‘Sinorhizobium adhaerens’ is not validly published. Opinion 84. Int J Syst Evol Microbiol 2008;58:1973–1973.

33. Lindstrom K, Young JPW. International Committee on Systematics of Prokaryotes; Subcommittee on the taxonomy of Agrobacterium and Rhizobium. Minutes of the meetings, 31 August 2008, Gent, Belgium. Int J Syst Evol Microbiol 2009;59:921–922.

34. Young JM. Sinorhizobium versus Ensifer: may a taxonomy subcommittee of the ICSP contradict the Judicial Commission? Int J Syst Evol Microbiol 2010;60:1711–1713.

35. Tindall BJ. The correct name of the taxon that contains the type strain of Rhodococcus equi. Int J Syst Evol Microbiol 2014;64:302–308.

36. de Lajudie P, Young JPW. International Committee on Systematics of Prokaryotes Subcommittee on the Taxonomy of Rhizobia and Agrobacteria Minutes of the meeting by video conference, 11 July 2018. Int J Syst Evol Microbiol 2019;69:1835–1840.

37. Martens M, Delaere M, Coopman R, De Vos P, Gillis M, et al. Multilocus sequence analysis of Ensifer and related taxa. Int J Syst Evol Microbiol 2007;57:489–503.

38. Kumar HKS, Gan HM, Tan MH, Eng WWH, Barton HA, et al. Genomic characterization of eight Ensifer strains isolated from pristine caves and a whole genome phylogeny of Ensifer (Sinorhizobium). J Genomics 2017;5:12–15.

39. Martens M, Dawyndt P, Coopman R, Gillis M, De Vos P, et al. Advantages of multilocus sequence analysis for taxonomic studies: a case study using 10 housekeeping genes in the genus Ensifer (including former Sinorhizobium). Int J Syst Evol Microbiol 2008;58:200–214.

40. diCenzo GC, Debiec K, Krzysztoforski J, Uhrynowski W, Mengoni A, et al. Genomic and biotechnological characterization of the heavy-metal resistant, arsenic-oxidizing bacterium Ensifer sp. M14. Genes 2018;9:379.

41. Garrido-Oter R, Nakano RT, Dombrowski N, Ma K-W, AgBiome Team, et al. Modular traits of the Rhizobiales root microbiota and their evolutionary relationship with symbiotic rhizobia. Cell Host Microbe 2018;24:155-167.e5.

42. Parks DH, Chuvochina M, Waite DW, Rinke C, Skarshewski A, et al. A standardized bacterial taxonomy based on genome phylogeny substantially revises the tree of life. Nat Biotechnol 2018;36:996–1004.

43. diCenzo GC, Cangioli L, Nicoud Q, Cheng JHT, Blow MJ, et al. DNA methylation patterns in bacteria of the genus Ensifer during free-living growth and during nitrogen-fixing symbiosis with Medicago spp. bioRxiv. Epub ahead of print 8 March 2021. DOI: 10.1101/2021.03.08.434416.

44. Wilks M. Predation Mediated Carbon Turnover in Nutrient-Limited Cave Environments. MSc Thesis; University of Akron; 2013.

45. Martin MO. Predatory prokaryotes: an emerging research opportunity. J Mol Microbiol Biotechnol 2002;4:467–477.

46. Wang ET, Tan ZY, Willems A, Fernández-López M, Reinhold-Hurek B, et al. Sinorhizobium morelense sp. nov., a Leucaena leucocephala-associated bacterium that is highly resistant to multiple antibiotics. Int J Syst Evol Microbiol 2002;52:1687–1693.

47. Wang YC, Wang F, Hou BC, Wang ET, Chen WF, et al. Proposal of Ensifer psoraleae sp. nov., Ensifer sesbaniae sp. nov., Ensifer morelense comb. nov. and Ensifer americanum comb. nov. Syst Appl Microbiol 2013;36:467–473.

48. Zweiger G, Marczynski G, Shapiro L. A Caulobacter DNA methyltransferase that functions only in the predivisional cell. J Mol Biol 1994;235:472–485.

49. Wright R, Stephens C, Shapiro L. The CcrM DNA methyltransferase is widespread in the alpha subdivision of proteobacteria, and its essential functions are conserved in Rhizobium meliloti and Caulobacter crescentus. J Bacteriol 1997;179:5869–5877.

50. Cohan FM. Towards a conceptual and operational union of bacterial systematics, ecology, and evolution. Phil Trans R Soc B 2006;361:1985–1996.

51. Parker CT, Tindall BJ, Garrity GM. International Code of Nomenclature of Prokaryotes. Int J Syst Evol Microbiol 2015;65:4284–4287.

52. de Lajudie P, Mousavi SA, Young JPW. International Committee on Systematics of Prokaryotes Subcommittee on the Taxonomy of Rhizobia and Agrobacteria. Minutes of the closed meeting by videoconference, 6 July 2020. Int J Syst Evol Microbiol 2021;71:004784.

53. Richter M, Rosselló-Móra R. Shifting the genomic gold standard for the prokaryotic species definition. Proc Natl Acad Sci USA 2009;106:19126–19131.

54. Meier-Kolthoff JP, Klenk H-P, Göker M. Taxonomic use of DNA G+C content and DNA– DNA hybridization in the genomic age. Int J Syst Evol Microbiol 2014;64:352–356.

55. Zhao J-J, Zhang J, Zhang R-J, Zhang C-W, Yin H-Q, et al. Rhizobium rhizosphaerae sp. nov., a novel species isolated from rice rhizosphere. Antonie van Leeuwenhoek 2017;110:651–656.

56. Peng G, Yuan Q, Li H, Zhang W, Tan Z. Rhizobium oryzae sp. nov., isolated from the wild rice Oryza alta. Int J Syst Evol Microbiol 2008;58:2158–2163.

57. Li Y, Yan J, Yu B, Wang ET, Li X, et al. Ensifer alkalisoli sp. nov. isolated from root nodules of Sesbania cannabina grown in saline–alkaline soils. Int J Syst Evol Microbiol 2016;66:5294–5300.

58. Merabet C, Martens M, Mahdhi M, Zakhia F, Sy A, et al. Multilocus sequence analysis of root nodule isolates from Lotus arabicus (Senegal), Lotus creticus, Argyrolobium uniflorum and Medicago sativa (Tunisia) and description of Ensifer numidicus sp. nov. and Ensifer garamanticus sp. nov. Int J Syst Evol Microbiol 2010;60:664–674.

59. Yan H, Yan J, Sui XH, Wang ET, Chen WX, et al. Ensifer glycinis sp. nov., a rhizobial species associated with species of the genus Glycine. Int J Syst Evol Microbiol 2016;66:2910–2916.

60. Nick G, de Lajudie P, Eardly BD, Suomalainen S, Paulin L, et al. Sinorhizobium arboris sp. nov. and Sinorhizobium kostiense sp. nov., isolated from leguminous trees in Sudan and Kenya. Int J Syst Evol Microbiol 1999;49:1359–1368.

61. Wei GH, Wang ET, Tan ZY, Zhu ME, Chen WX. Rhizobium indigoferae sp. nov. and Sinorhizobium kummerowiae sp. nov., respectively isolated from Indigofera spp. and Kummerowia stipulacea. Int J Syst Evol Microbiol 2002;52:2231–2239.

62. Rome S, Fernandez MP, Brunel B, Normand P, Cleyet-Marel JC. Sinorhizobium medicae sp. nov., isolated from annual Medicago spp. Int J Syst Bacteriol 1996;46:972–980.

63. Lloret L, Ormeño-Orrillo E, Rincón R, Martínez-Romero J, Rogel-Hernández MA, et al. Ensifer mexicanus sp. nov. a new species nodulating Acacia angustissima (Mill.) Kuntze in Mexico. Syst Appl Microbiol 2007;30:280–290.

64. Chen W, Sheng X-F, He L-Y, Huang Z. Rhizobium yantingense sp. nov., a mineral-weathering bacterium. Int J Syst Evol Microbiol 2015;65:412–417.

65. Zhang X, Li B, Wang H, Sui X, Ma X, et al. Rhizobium petrolearium sp. nov., isolated from oil-contaminated soil. Int J Syst Evol Microbiol 2012;62:1871–1876.

66. Zhang S, Yang S, Chen W, Chen Y, Zhang M, et al. Rhizobium arenae sp. nov., isolated from the sand of Desert Mu Us, China. Int J Syst Evol Microbiol 2017;67:2098–2103.

67. Turdahon M, Osman G, Hamdun M, Yusuf K, Abdurehim Z, et al. Rhizobium tarimense sp. nov., isolated from soil in the ancient Khiyik River. Int J Syst Evol Microbiol 2013;63:2424–2429.

